# SENP2-based N-terminal truncation of α-synuclein in Lewy pathology propagation

**DOI:** 10.1101/2024.09.18.613608

**Authors:** Katsutoshi Taguchi, Yoshihisa Watanabe, Masaki Tanaka

## Abstract

α-Synuclein (αSyn) is a major component of Lewy bodies (LBs) and Lewy neurites (LNs) which are pathological features of Parkinson’s disease (PD) and Dementia with Lewy bodies. In the PD brain, with disease progression, LB/LN formation is propagated from the lower brainstem to the cerebral cortex. Prion-like cell-to-cell seed-transmission has been implicated as an underlying mechanism for Lewy-pathology propagation. However, the biochemical properties and production mechanism of those pathogenic seeds are unelucidated. In this study, we ascertained that the seeds released from pathological neurons that harbour LB/LN-like aggregates have the N-terminally truncated form of αSyn. This N-terminal truncation is directly catalysed by SENP2, which is a well-known deSUMOylation enzyme. After SENP2 processing of recombinant αSyn, the SDS-resistant high-molecular oligomer formation was promoted *in vitro*. Inhibition of SENP2 activity suppressed aggregate formation and propagation in cultured neurons and mouse brains. Thus, SENP2 might be a novel therapeutic target in LB diseases.

## Introduction

α-Synuclein (αSyn) is one of the major constituents of Lewy bodies (LBs) and Lewy neurites (LNs) that constitute the well-known pathological hallmarks of synucleinopathies, including Parkinson’s disease (PD) and dementia with Lewy bodies (DLB)^1–3^. In the sporadic PD brain, the propagation of LB/LN formation indicates a caudorostral process that is associated with disease progression from the lower brainstem, through the basal midbrain and forebrain, into the cerebral cortex^4,5^. Interestingly, LBs/LNs were observed even in the grafted dopaminergic neurons derived from foetal mesencephalic tissue that has been transplanted into the striatum of a patient with PD^6^. This suggests the presence of prion-like propagation as the underlying mechanism in neurodegeneration^7^. In several experimental models, inoculation of preformed fibrils (PFFs) prepared from recombinant αSyn or PD brain-derived tissue extracts into the mouse striatum can induce LB/LN-like aggregate formation and propagation through the anatomical connections in the synaptic contacts within the brain^8–10^.

Abnormal intracellular aggregates, such as LBs and LNs, are formed by the recruitment of intrinsic soluble αSyn into the insoluble aggregate core (the so-called “seeds”)^7,11^. Therefore, the endogenous expression of αSyn as well as seeds, which accelerate the polymerization of soluble αSyn, are required for the formation of aggregates. Next, the cell-to-cell transmission of seeds is a crucial step for the propagation of Lewy pathology. However, the biochemical properties and mechanism of production of these pathogenic seeds that are released from pathological neurons, which harbour LBs/LNs, are unelucidated.

The evidence from recent studies implicates posttranslational modifications (PTMs) of αSyn, such as phosphorylation, glycosylation, acetylation, ubiquitination, SUMOylation, and truncation, in the pathophysiological propensities of αSyn^12,13^. By using histological methods and mass spectrometric analysis, the presence of various PTMs of αSyn has been detected in the PD brain. Phosphorylation at the Ser129 residue is a well-known PTM of αSyn-comprising LBs/LNs and is used as a marker for the detection of these abnormal aggregates^14^. Furthermore, ubiquitination and SUMOylation play important roles in the regulation of αSyn’s stability and aggregation ability^15,16^. The truncation of αSyn has significant effects on the aggregation ability, cross-seeding kinetics, and fibril stability of the resultant species^12,17,18^. Interestingly, there are some characteristic differences between the C-terminal- and N-terminal-truncated forms of the αSyn^17,19^, wherein the former shows higher aggregation ability and the latter shows lower aggregation ability and fibril stability with higher cross-seeding activity in cultured cells and the mouse brain. The injection of PFFs prepared from the N-terminal 10-residue- or 30-residue-truncated αSyn, rather than PFFs prepared from full-length αSyn, into the mouse striatum induces more abundant aggregate formation^17^, which, given the consensus on the inverse correlation of seeding activity with fibril stability in the field of prions, possibly contributes to the lower stability of N-terminal-truncated αSyn-derived PFFs^20^. Among the various αSyn-processing enzymes that have been identified thus far, caspase-1 and asparagine endopeptidase are C-terminal cleavage enzymes, whereas 20S proteasome, calpain-1, cathepsins, and matrix metalloproteases can truncate both C-terminal and N-terminal regions, including the NAC region of αSyn^12^. Furthermore, PFFs are processed by lysosomal cathepsin B, and this partial cleavage enhances the seeding ability of PFFs^21^.

Studies using mass spectrometric analysis have identified many cleavage sites within the tissue extracts obtained from the PD brain^12^. For example, calpain-mediated C-terminal truncation at 122/123 residues of αSyn has been detected in LBs/LNs^22,23^; however, the relationship between αSyn truncation and the biochemical properties of the seeds remains elusive.

In this study, we explored this relationship through the characterization of seed-constituting αSyn released from PD-model neurons containing pathological αSyn aggregates and identified novel cleavage sites in the N-terminal region.

## Results

### Isolation and characterization of cell-to-cell transmissible seeds

To isolate the cell-to-cell transmissible seeds released from pathological neurons that harbour LB/LN-like abnormal aggregates, the oligomeric species of αSyn within the conditioned medium of those pathological neurons were screened using sucrose density-gradient ultracentrifugation and Western blotting analysis (Figure S1A–C and Methods). Accordingly, 140-kDa αSyn oligomers, separated by native-PAGE, were recovered from the interface fraction between the concentrated conditioned medium and the 0.8 M sucrose-containing PBS layer. Moreover, we confirmed that the LB/LN-like aggregate formation in the primary cultured neurons was induced by the fraction-containing fresh medium (Figure S1D).

To more precisely isolate the seeds and obviate PFF contamination, isolation was achieved with microfluidic devices (Figure 1A, B; C1 and C3 chambers), wherein seeds released from pathological neurons were recovered from the PFF-free culture medium. Furthermore, using this microfluidic method, 140-kDa αSyn oligomers that were separated by native-PAGE were recovered from the interface fraction as well as the results obtained by the batch method (Figure 1C–E). Atomic force microscopy showed that this fraction contained a uniform population of granular molecules with a diameter of 36 ± 1.2 nm (Figure 1F, G). Next, to further confirm that these granular molecules are composed of αSyn, these molecules were detected by several antibodies against αSyn. For αSyn detection by immunoelectron microscopy, the reactivity of secondary antibodies was optimized using PFFs as the positive control (Figure S1G). These granular molecules (diameter 35 ± 2.3 nm) detected by electron microscopy were composed of αSyn and thus labelled by certain monoclonal antibody 5G4, but not by the O2 antibody (Figure 1H–J). 5G4 recognizes protofibrils or large oligomeric species of αSyn^28^. To determine the molecular weight of the intact seed structure, size-exclusion chromatography (SEC)-high-performance liquid chromatography (HPLC) analysis was performed (Figure S1E, F). The seed structure detected with immunoelectron microscopy was recovered in the SEC Fraction No. 8 (corresponding to approximately 1000 kDa), as determined using a high-molecular-weight standard kit. These results indicated that seeds of approximately 1000-kDa size were produced in the pathological neurons and subsequently released into the culture medium.

**Figure 1.**
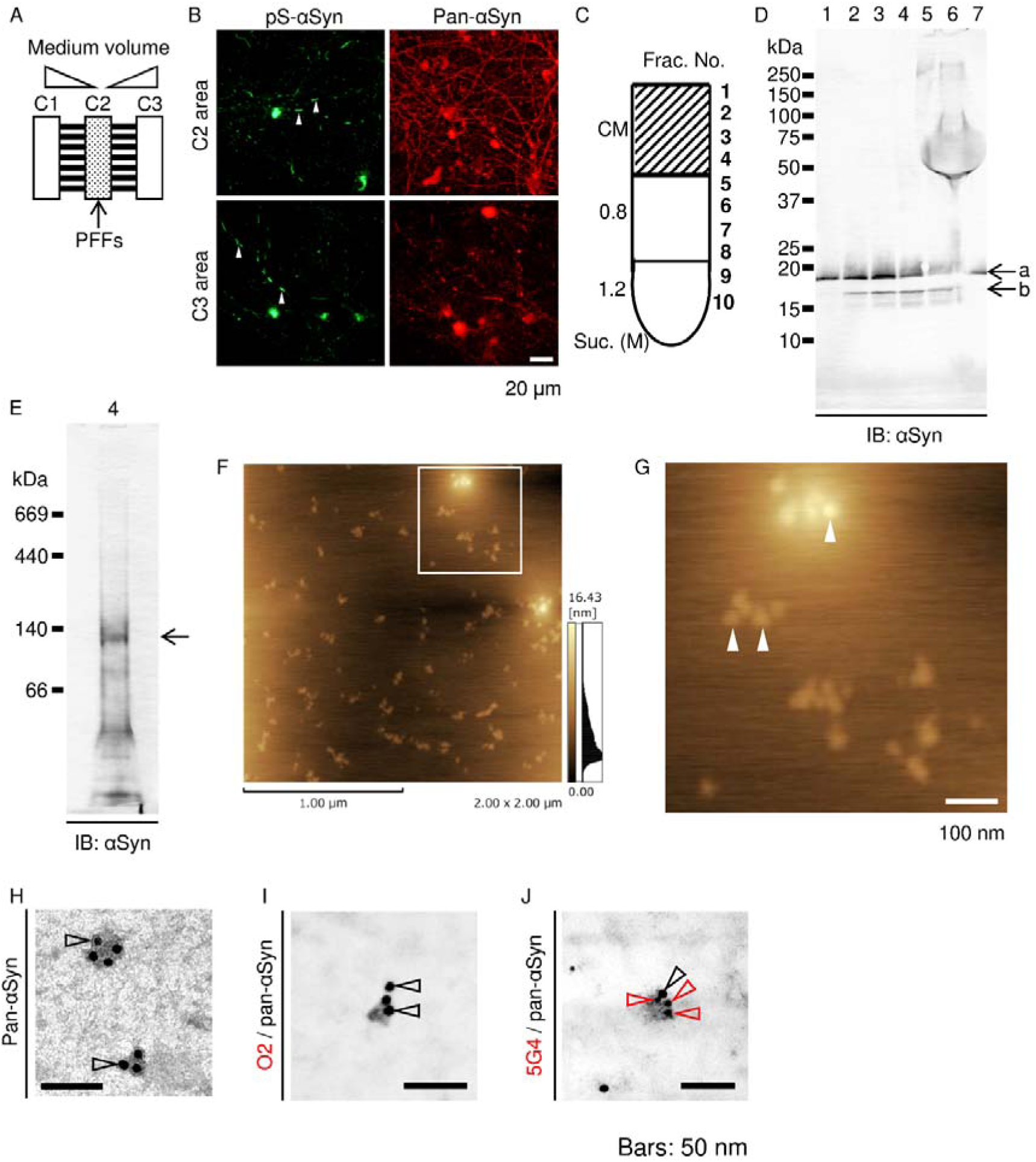
Isolation and characterization of pathological seeds. **(A)** Schematic of a microfluidic device for pathological seed preparation. The primary neurons were cultured in separate chambers (C1, C2, and C3) connected through microgrooves. Preformed fibrils (PFFs) were supplemented in C2 culture medium, with height kept constantly lower than that of C1 and C3 media during cell culture. **(B)** Propagation of phosphorylated-αSyn (pS-αSyn)-positive intracellular aggregates from C2 (upper panels) to C1 and C3 (lower panels) areas. **(C)** Fractionation of concentrated culture medium (CM) recovered from C1 and C3 using sucrose density-gradient ultracentrifugation. **(D)** SDS-PAGE and immunoblotting assay using anti-αSyn antibody (numbers indicate recovered fractions in **(C)**; His-tagged and endogenous αSyn (arrows a and b, respectively)). **(E)** Native-PAGE and immunoblotting assay of Fraction 4 **(C, D)**. A major band at approximately 140-kDa (arrow) was detected using anti-αSyn antibody. **(F, G)** Atomic force microscopic imaging of the molecules recovered in Fraction 4, containing uniform granular molecules (diameter 36 ± 1.2-nm; n=16). White square in **(F)** is magnified in **(G)**. Arrowheads in **(G)** indicate the examples of those molecules. **(H–J)** Immunoelectron microscopic imaging of molecules in Fraction 4; αSyn-positive granular molecules detected using anti-pan-αSyn antibody (black arrowheads: 10-nm gold particles) in **(H–J)**. Granular molecules were recognized by anti-aggregated αSyn antibody 5G4 (**J**; red arrowheads, 6-nm gold particles), but not by the anti-oligomeric αSyn antibody O2 (**I**; negative for 6-nm gold particles). αSyn-positive granular molecules (diameter 35 ± 2.3 nm; n=6). Data represent mean ± SEM. Scale bars: 20 μm and 1 μm in **B** and **F**, respectively; 100 nm in **G**, and 50 nm in **H–J**.

### Seeds contain truncated forms of αSyn

To investigate whether seeds contained the truncated αSyn or not, the cleavage sites within αSyn were screened using mass spectrometric analysis. As a large amount of seed protein was required for identification of the cleavage sites, a large microfluidic device was developed (Figure S2). Using immunoprecipitation, samples for mass spectrometric analysis were prepared from seed fraction (Figure 2A). PFF-constituting αSyn, which was derived from the human αSyn sequence, was prepared as a control sample for mass spectrometric analysis because results of the artificially cleaved sites detected in PFF-constituting αSyn were subtracted from those of the cleaved sites detected in the seed-constituting αSyn. Protein bands, Nos. 1–5, and control bands were subjected to mass spectrometric analysis (Figure 2B), and the results suggested the presence of 6 cleavage sites (Figure 2C). The C-terminal cleavage site at the 119/120 residue is a 20S proteasome-processing site^24^. To our knowledge, cleavage sites detected at the N-terminal and NAC regions have not been reported previously. Interestingly, conserved cleavage sites (KX|KXGV), the N-terminal sides of lysine residues located at the N-terminal repeat regions (1–4) of αSyn, were detected (Figure 2C). Although the cleavage site at repeat 4 was detected in the results of the control sample in the mass spectrometric analysis, this site was considered as a possible cleavage site, because the primary amino acid sequence is completely conserved at repeat 3. These results suggested that the N-terminus of the seed-constituting αSyn was partially processed by unknown N-terminal lysine endopeptidase activity. This information on the conserved processing sites is useful for identifying the processing enzyme. Therefore, we focused on these N-terminal cleavage sites of αSyn.

**Figure 2.**
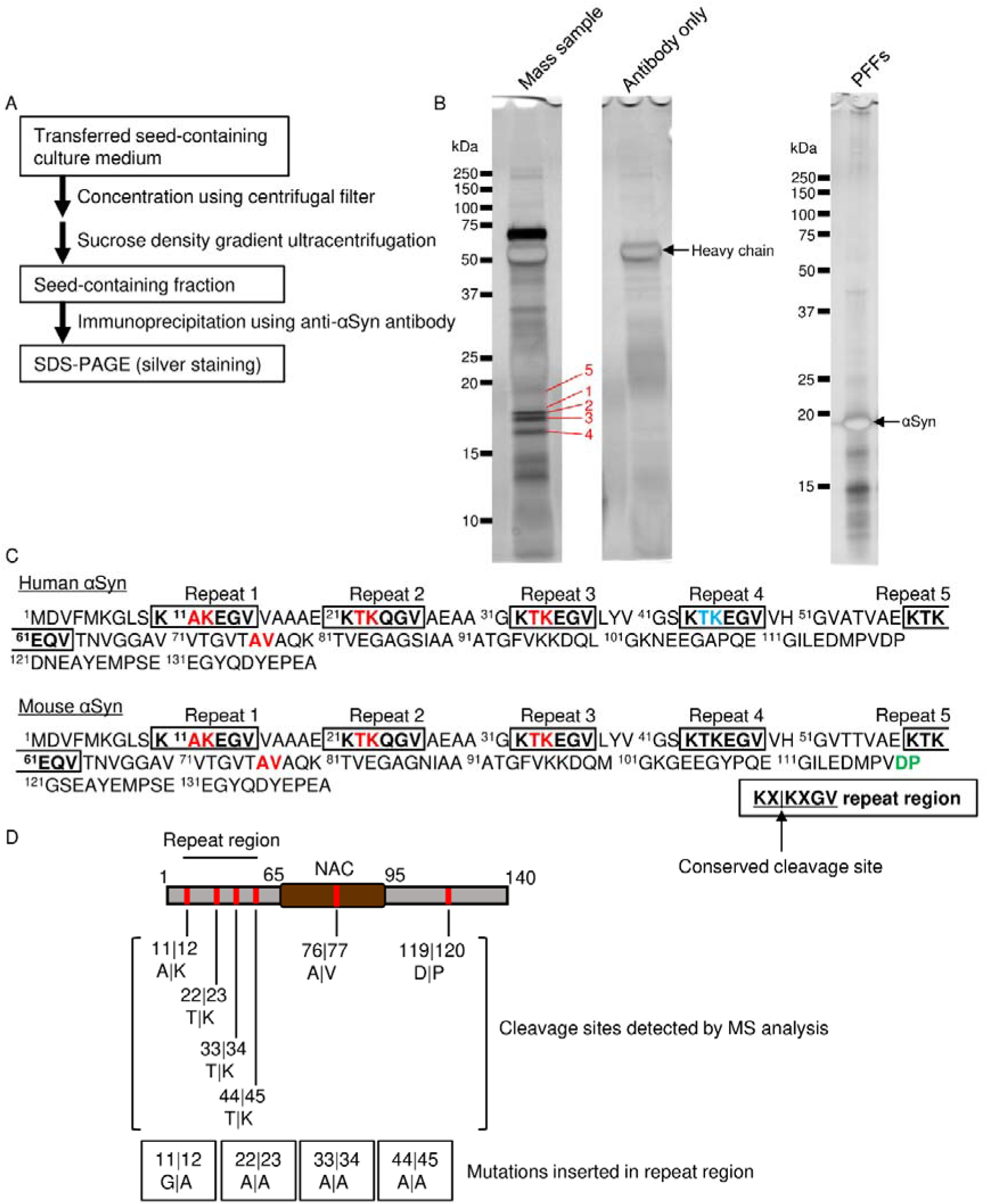
Truncated αSyns are included in the pathological seeds. **(A)** Experimental design to prepare samples for the detection of αSyn truncation using mass spectrometric analysis. **(B, C)** Immunoprecipitated proteins were separated by SDS-PAGE and detected by silver staining as in **(B)**. Gel bands indicated by the numbers 1–5 were dissected and subjected to mass spectrometric analysis. Predicted truncation sites are shown in **(C)**. Truncated sites detected in common amino acid sequence between human and mouse αSyn are indicated in red, whereas sites detected in human or mouse αSyn-specific sequences are indicated in blue or green, respectively. **(D)** Schematic representation of the predicted cleavage sites and the corresponding insertion of each mutation to inhibit the cleavage.

### αSyn is processed by SENP2

Based on the results of mass spectrometric analysis, candidate enzymes that cleave the N-terminal side of lysine residues located at the N-terminal repeat regions of αSyn (conserved cleavage sites in Figure 2C) were screened using the Peptidase database, MEROPS, which showed that carboxypeptidase B, USP7, and SENP2 were candidate enzymes that can process the N-terminal side of the lysine residues. After screening these enzymes, each enzyme’s inhibitors were selected (Table 1). To examine the inhibitory effect of the inhibitors on aggregate formation, primary neurons were cultured and treated with PFFs in the presence of each inhibitor. In the inhibitor assay, NSC632839 could significantly inhibit the aggregate formation (Figure 3A–F), whereas the reagent inhibits the enzymatic activity of not only USP7 and SENP2 but also USP2 and UCHL1^25,26^. Therefore, to examine whether αSyn is directly processed by these enzymes, protease assays were performed using recombinant active enzymes. Recombinant αSyn was cleaved by SENP2, and the resulting fragment was detected clearly (Figure 3G). However, cleaved fragments of αSyn were not detected in the presence of USP7, USP2, and UCHL1 (Figure S4A–C). SENP2 preferentially cleaved the soluble form of αSyn, rather than PFFs. The cleaved fragment of αSyn increased, depending on the SENP2 dose (Figure 3H, I). Interestingly, SDS-resistant high-molecular-weight signals of αSyn were increased by high-dose SENP2 treatment. This cleavage pattern of αSyn by SENP2 was diminished by the insertion of mutations at the predicted cleavage sites, as represented in Figure 2D (Figure 3J).

**Figure 3.**
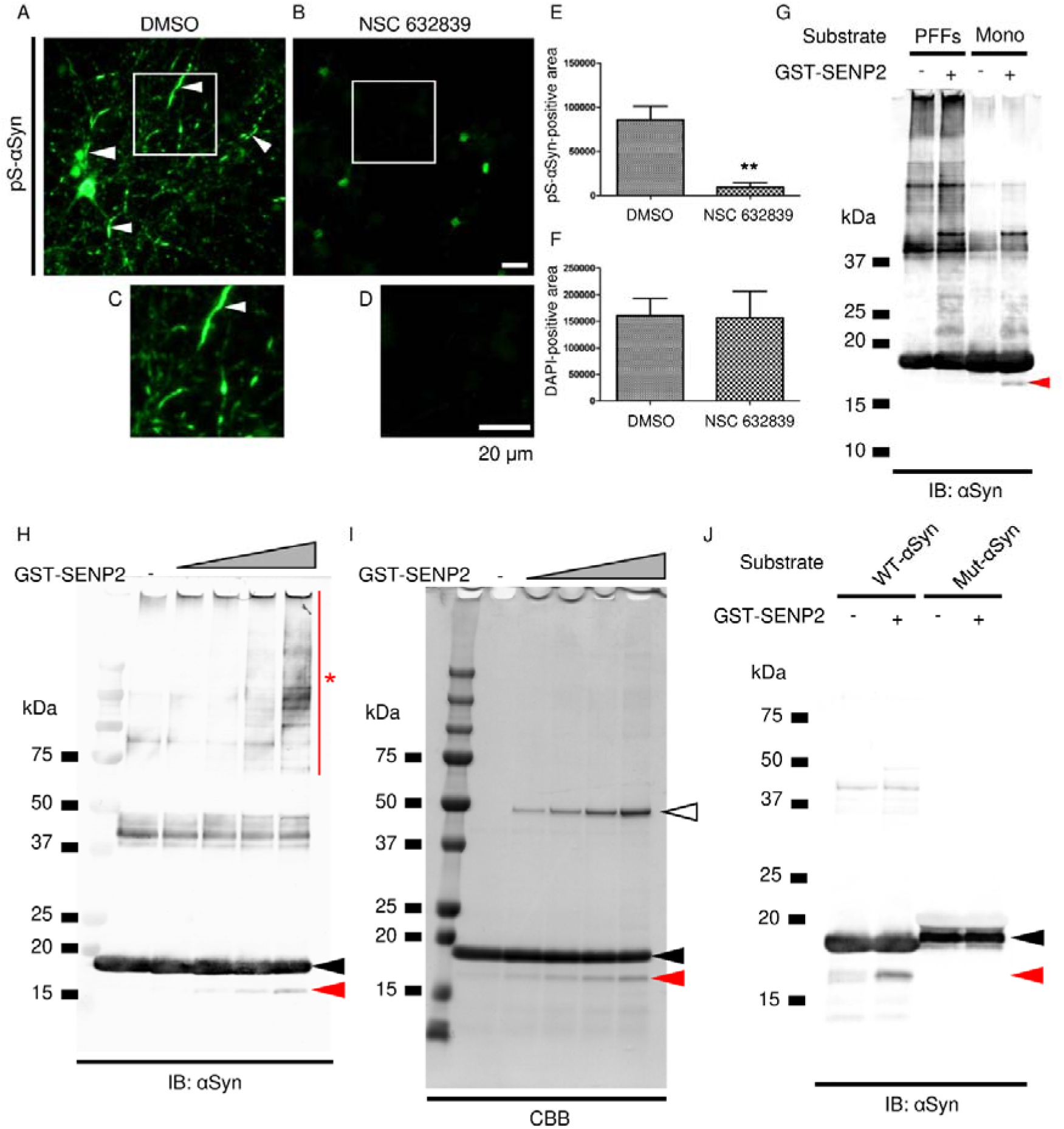
Identification of processing enzyme. **(A–F)** Inhibition of aggregate formation by NSC632839. Primary neurons were treated with PFFs and cultured in the presence of DMSO **(A)** or NSC632839 **(B)**. The region marked by a white square in **(A)** and **(B)** is magnified in **(C)** and **(D)**, respectively. Arrowheads in **(A)** and **(C)** indicate the formation of the aggregates. Quantitative comparison of the phosphorylated-αSyn-positive area **(E)** and DAPI-positive area **(F)** between DMSO-treated and NSC632839-treated neurons (n=3 fluorescent images from 2 coverslips were analysed for each treatment; \*\**p* < 0.01, two-tailed, unpaired *t*-test; two independent cultures were performed and confirmed reproducibility). **(G)** Cleavage of αSyn by GST-fused recombinant SENP2 for 4 h at 37°C. A cleaved fragment is indicated by the arrowhead. Mono, αSyn monomer. **(H–J)** Dose-dependent effect of SENP2 on αSyn-cleavage. αSyn was processed for 24 h at 37°C. The cleaved fragment (red arrowhead) was gradually increased in the presence of a higher amount of SENP2. Black and white arrowheads show His-tagged full-length αSyn- and GST-fused SENP2 recombinant protein, respectively. Note that αSyn-cleavage increased high-molecular-weight SDS-resistant immunoreactivity (asterisk). This αSyn-cleavage by SENP2 for 4 h was inhibited by the insertion of the N-terminal mutations **(J)**. Scale bar: 20 μm in **(A–D)**.

**Table 1.**
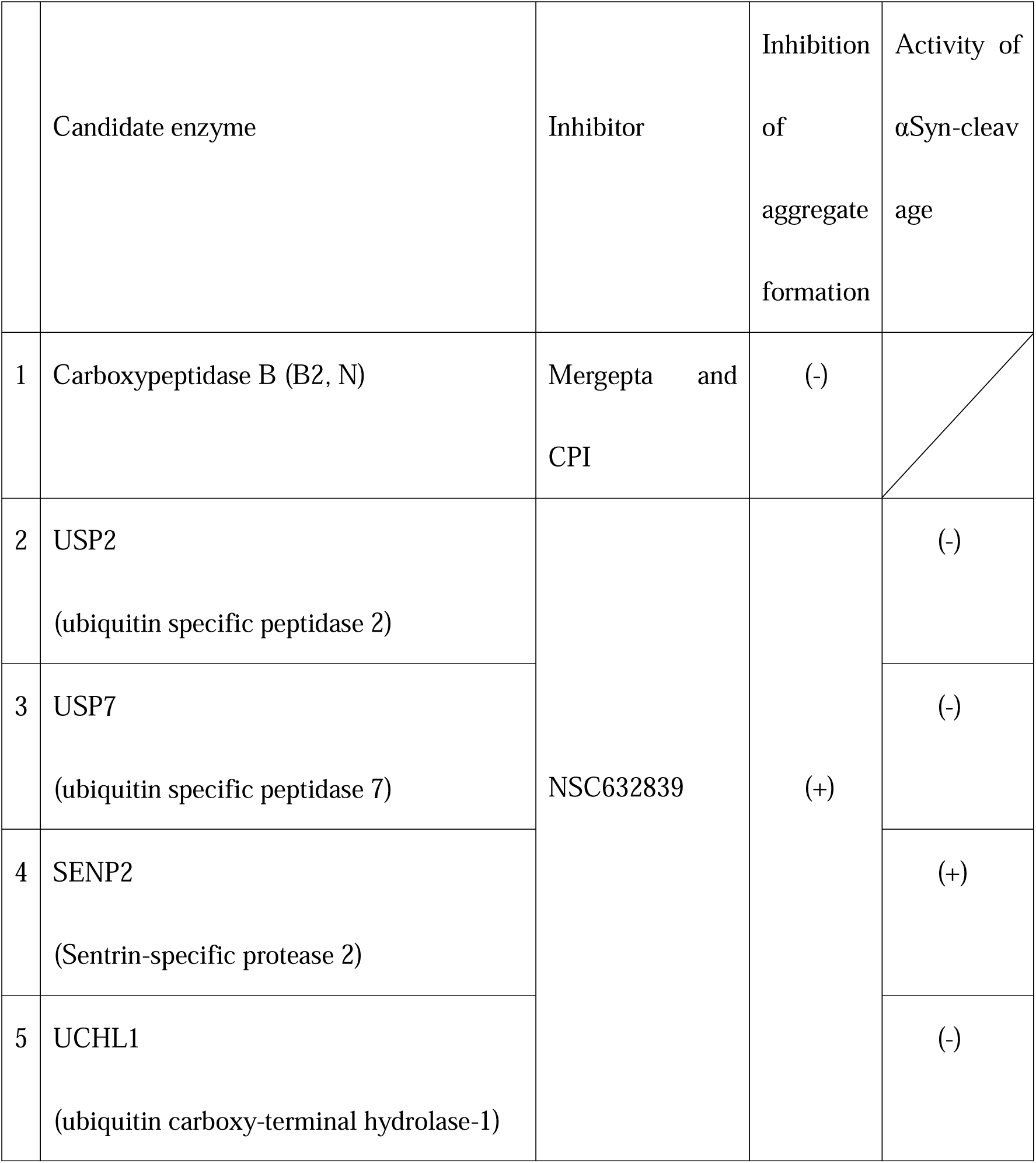
Screening of αSyn-processing enzyme.

SENP2 was ordinarily expressed in the nucleus^27^. After aggregate formation, SENP2 was partially distributed in the aggregates (Figure 4). In cultured neurons harbouring phosphorylated-αSyn aggregates induced by PFFs, SENP2 was partially detected in those aggregates (Figure 4A). In the human PD brain, SENP2 was observed in the amorphous aggregates composed of phosphorylated αSyn, but not in typical LBs, which are spheroidal (Figure 4B and Figure S5C, D).

**Figure 4.**
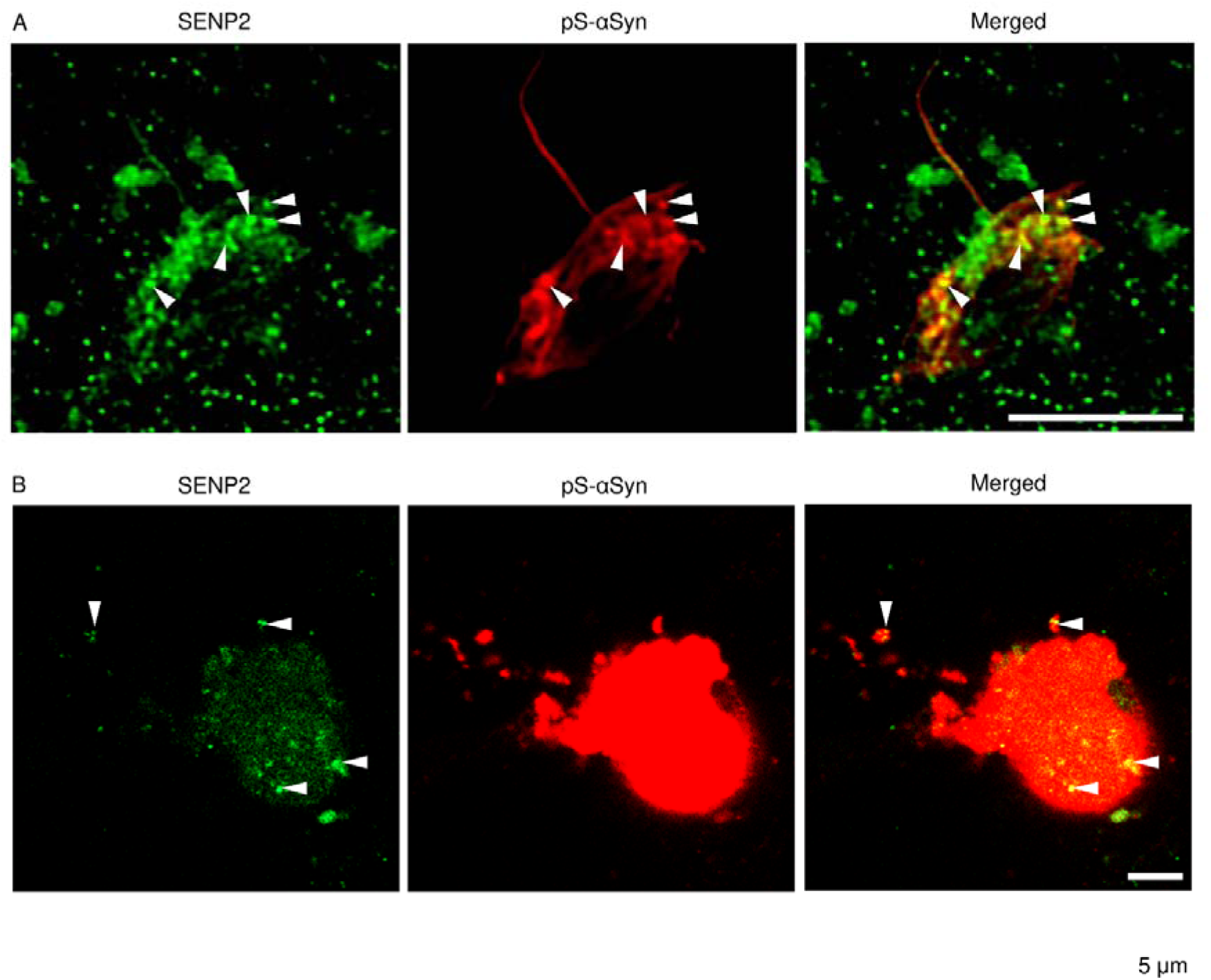
Presence of endogenous SENP2 in αSyn-aggregates. **(A)** Presence of SENP2 in PFF-induced αSyn-aggregates formed in primary cultured neurons. **(B)** Presence of SENP2 in αSyn-aggregates formed in the neuron of the substantia nigra pars compacta of PD-brain. Arrowheads indicate examples of the overlapping intense signals of SENP2 and phosphorylated αSyn. Scale bar: 5 μm.

### Inhibition of SENP2 suppresses aggregate formation and its propagation

Biochemical studies demonstrated that SENP2 cleaved the N-terminal repeat regions of αSyn whereas its inhibitor, NSC632839, suppressed aggregate formation in cultured neurons. Next, we studied the inhibitory effects of SENP2 through the inhibitor infusion on aggregate formation and its propagation *in vivo*. The efficiency of the aggregate formation and inhibitory effect of NSC632839 on the propagation in the brain was examined using immunohistochemical methods. Injection of PFFs into the striatum (Figure 5A) showed aggregate formation, which was propagated from the striatum to other brain regions, such as the piriform cortex and amygdala, whereas inhibitor infusion into the lateral ventricle decreased aggregate formation and its propagation (Figure 5B). The number of cell bodies harbouring phosphorylated-αSyn-positive aggregates was significantly decreased in the piriform cortex and amygdala (Figure 5C, D). In this process, endogenous expression of soluble αSyn was required for aggregate formation, because αSyn-KO mice did not show any spreading of phosphorylated αSyn aggregates after the PFF injection (Figure S6). These results indicated that the inhibition of the enzymatic activity attenuated aggregate formation and its propagation efficiency.

**Figure 5.**
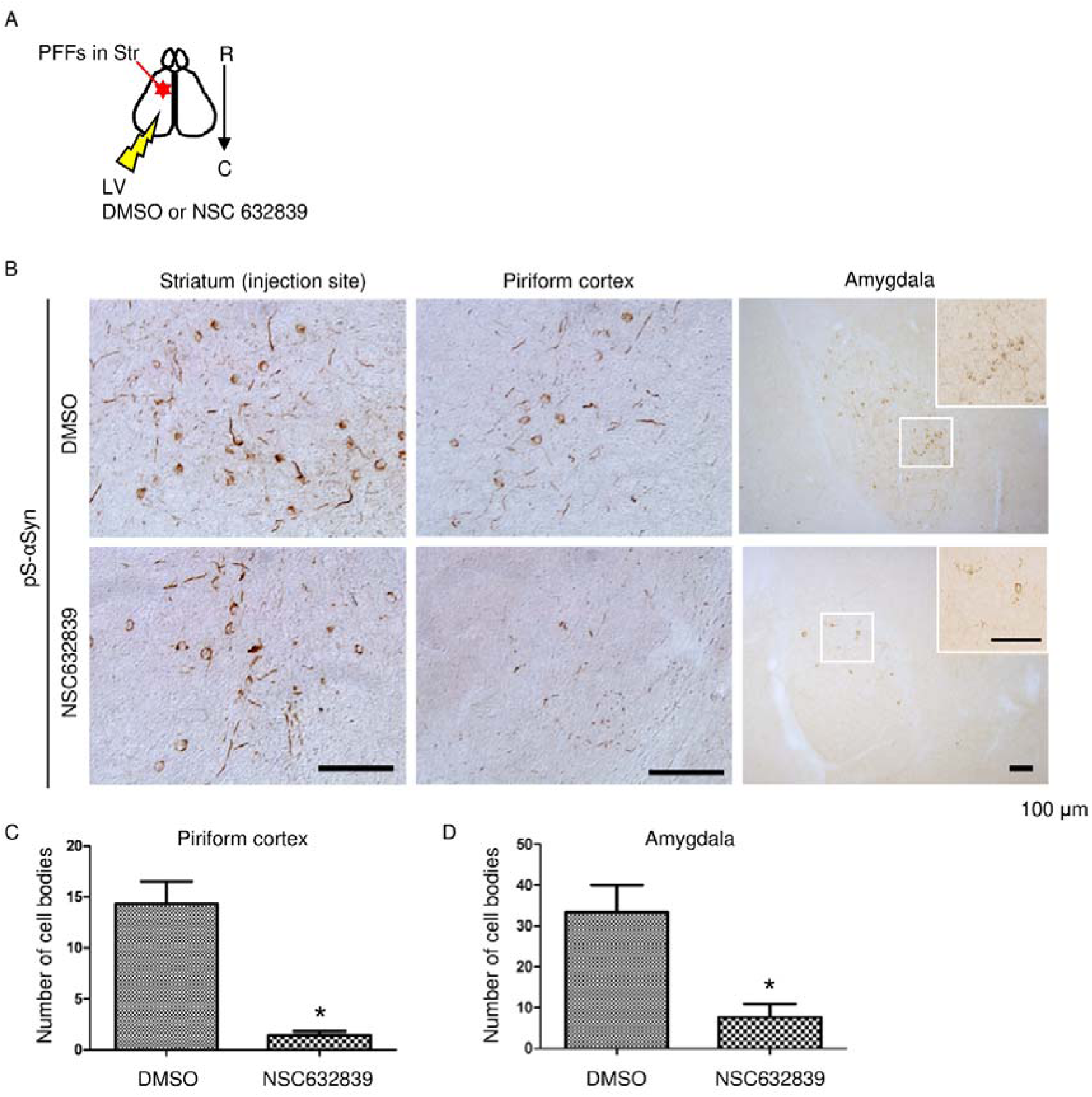
Inhibition of aggregate formation and cell-to-cell seed transmission. **(A)** Schematic representation of PFF injection in the WT mouse striatum, and DMSO or NSC632839 osmotic pump-based infusion into the lateral ventricle. C, caudal; LV, lateral ventricle; R, rostral; Str, striatum. **(B)** Inhibitory effect of NSC632839 infusion on the propagation of αSyn-aggregate formation in the ipsilateral sites of the striatum (injection site), piriform cortex, and amygdala. The region marked by a white square in the amygdala is magnified in each treatment. **(C, D)** Statistical analysis of the number of cell bodies harbouring pathological aggregates of phosphorylated αSyn, with or without NSC632839 treatment (**C**, n=5 NSC632839-treated and n=3 DMSO-treated control mice, the ipsilateral piriform cortex was analysed by using the two-tailed, unpaired *t*-test with Welch’s correction; **D**, n=3 mice in each treatment for the ipsilateral amygdala analysed by the two-tailed, unpaired *t*-test; \**p* < 0.05). Bar: 100 μm in **(B)**.

## Discussion

Recent studies indicated that cell-to-cell seed transmission is a pivotal mechanism underlying the propagation of Lewy-pathology in the PD brain^7^; however, the properties of seeds remain unelucidated. In this study, SEC-HPLC analysis demonstrated that seeds released from pathological cultured neurons were recovered as αSyn oligomers (approximately 1000 kDa), whereas native-PAGE assay showed a 140-kDa band. These results suggest that intact seeds are composed of 140-kDa rigid oligomers as the elementary units. During electrophoresis, the recovered seeds might become dissociated from the original high-molecular structure into those elementary units. These seeds were preferentially recognized by monoclonal antibody 5G4, but not O2. Furthermore, LBs and LNs were recognized by these antibodies^28,29^. Although LBs/LNs have epitopes against both 5G4 and O2 on their surfaces, seeds have only 5G4 epitopes on their molecular surfaces. The different structural milieu between LBs/LNs and the seeds might be responsible for this distinct immunoreactivity.

Biochemical and mass spectrometric analysis showed that seeds contained truncated forms of αSyn. Interestingly, N-terminal sides of lysine residues located in the N-terminal repeat regions of αSyn were processed by SENP2 – an enzyme that routinely removes SUMO from various SUMOylated proteins^30^. However, in the present study, SENP2 directly cleaved recombinant αSyn itself. After αSyn cleavage by high-dose SENP2, the formation of SDS-resistant high-molecular oligomeric species was facilitated by this treatment (see, Figure 3H, I). Furthermore, this enzymatic activity was crucial for aggregate formation. The inhibitory effects of NSC632839 on aggregate formation and its propagation were significantly observable *in vitro* and *in vivo*. Therefore, we further investigated the knockdown effects of SENP2 using AAV-assisted shRNA expression system in primary cultured neurons. However, SENP2 knockdown severely impaired normal neuronal development. Moreover, neuronal viability was decreased by the knockdown treatment during the neuronal maturation. On the other hand, significant knockdown efficiency was not obtained using matured neurons. These results suggested that SENP2 was indispensable for neuronal development, and its protein turnover rate in immature neurons might be higher than that in matured neurons. In the present study, NSC632839-based inhibition of SENP2 did not induce cellular damage, and this might be related to the neuronal maturation level.

Previous studies indicated that N-terminal repeat regions of αSyn implicate lipid membrane association with conformational changes to α-helical structure^31,32^. N-terminal truncation causes β-sheet structure-enriched conversion within the truncated αSyn^31^ and the resultant decreased stability of PFFs^17^. Thus, the N-terminal truncation of αSyn might promote pathogenesis, including αSyn oligomerization, and further provide an intracellular milieu that readily accelerates seed production.

Interestingly, a recent study revealed the presence of various kinds of αSyn-PTMs, such as acetylation, methylation, and carboxymethylation, in tissue extracts prepared from Lewy body disease (LBD) and control brains^33^. According to this study, N-terminal lysine residues at K12, K23, K34, and K45, which are SENP2-processing sites, are post-translationally modified with acetylation. The seeding ability of soluble αSyn harbouring K45Q, which is one of the acetylation mimic mutations, is significantly decreased. On the other hand, lysine residues at K12 and K23 are modified with carboxymethylation in LBD brains, but not in control brains. The seeding ability of αSyn might be regulated by the balance of PTMs, including truncation, acetylation, and carboxymethylation.

SENP2 was partially distributed in aggregates of phosphorylated αSyn in cultured neurons and PD brain. However, that was not detected in a typical LB, which might be attributable to the maturation stages of the aggregates. In the PD brain, SENP2 was detected in amorphous aggregates, rather than spheroidal aggregates, such as typical LBs, which are larger and more rigid than amorphous aggregates. SENP2 cleaves soluble αSyn, and the resulting fragments mediate aggregate formation. SENP2 may persist in immature aggregates but might be excluded from matured aggregates.

## Conclusion

Transmissible seeds contained the N-terminal truncated αSyn. SENP2 directly processed N-terminal repeat regions of αSyn, and inhibition of its activity suppressed aggregate formation and propagation in cultured neurons and mouse brains. The present study revealed that SENP2 activity is essential for Lewy-pathology formation. Thus, SENP2 constitutes a potential novel therapeutic target, and fine-tuning the PTMs of αSyn, including N-terminal truncation, could provide an innovative therapeutic strategy to prevent the progression of LB diseases.

## Limitations of this study

The presence of the characteristic seeds and the N-terminal processing sites of αSyn in the human brain are unconfirmed. Due to the limited amount of seeds recovered, it is currently challenging to produce structure-specific antibodies for *in vivo* seed detection using microfluidic devices. Further studies are required for precise seed characterization.

## STAR□Methods

### Key resources table

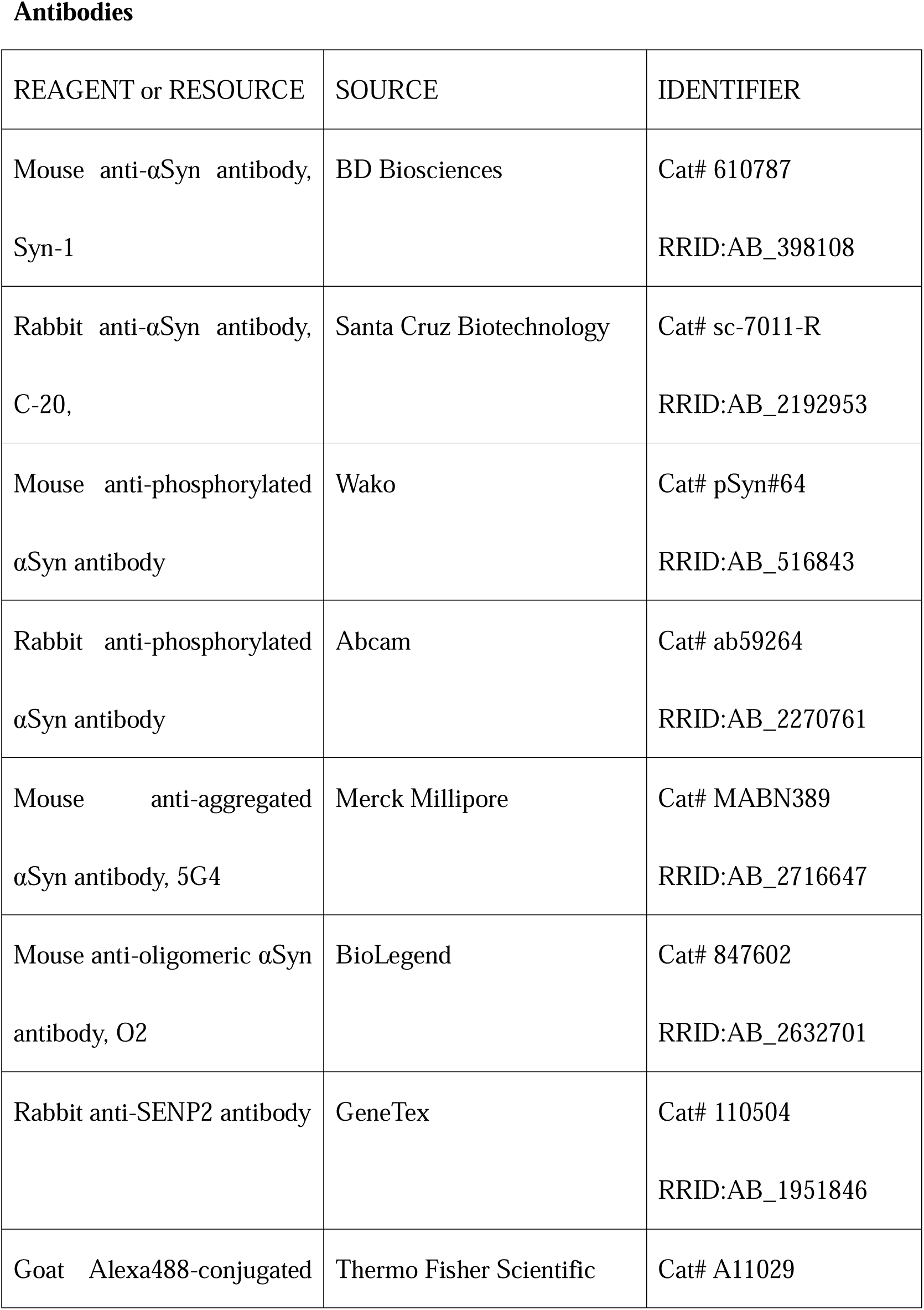
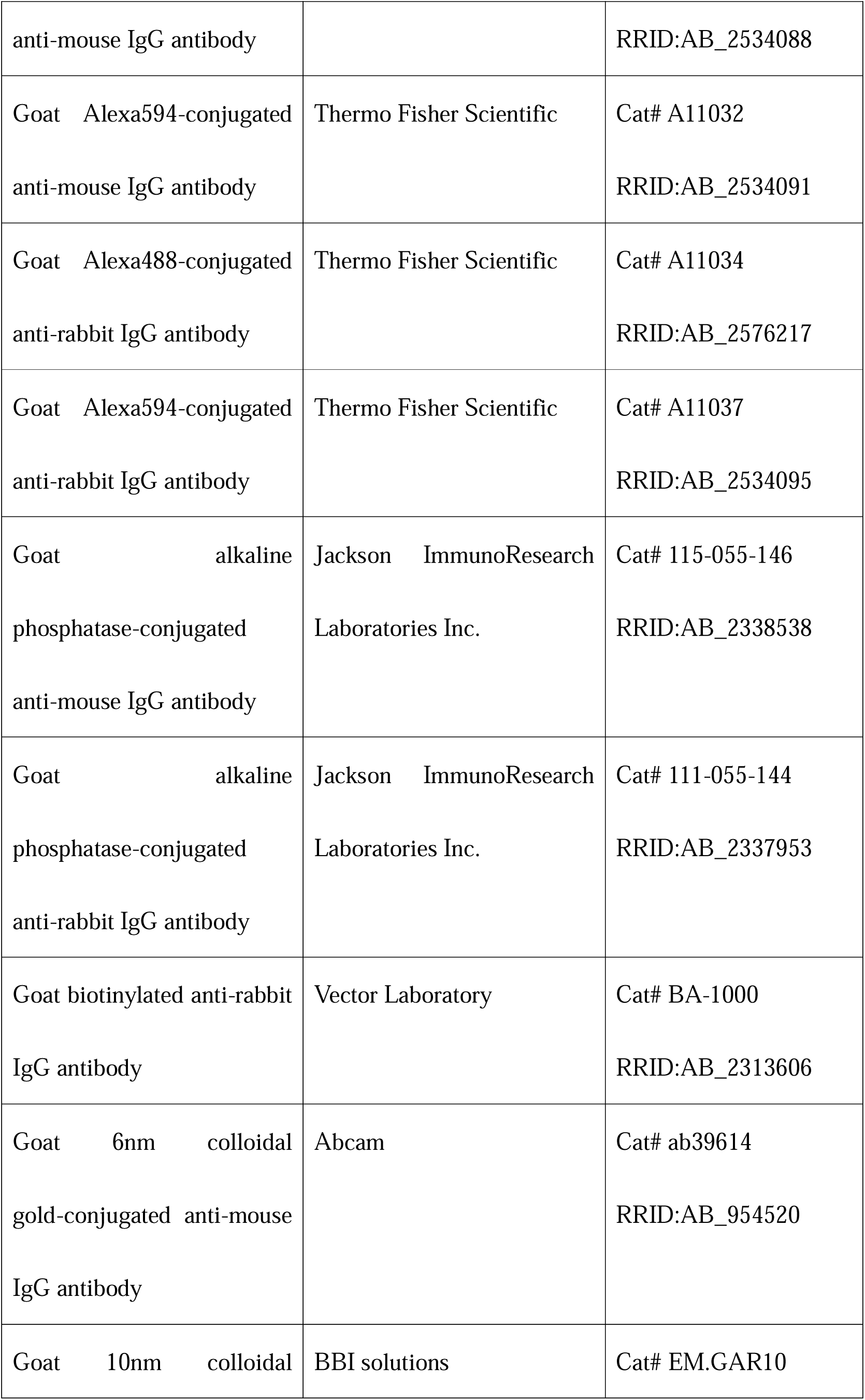
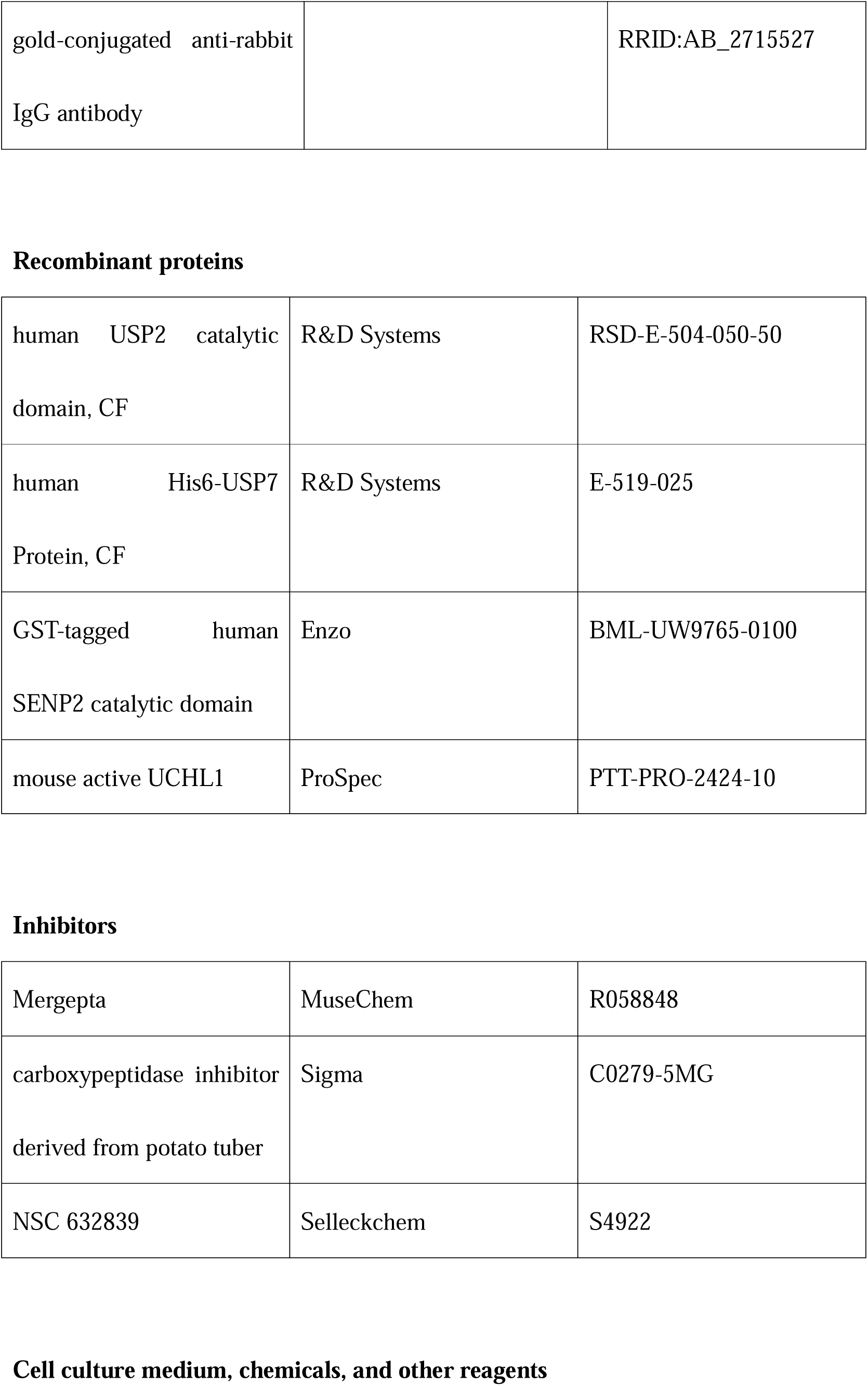
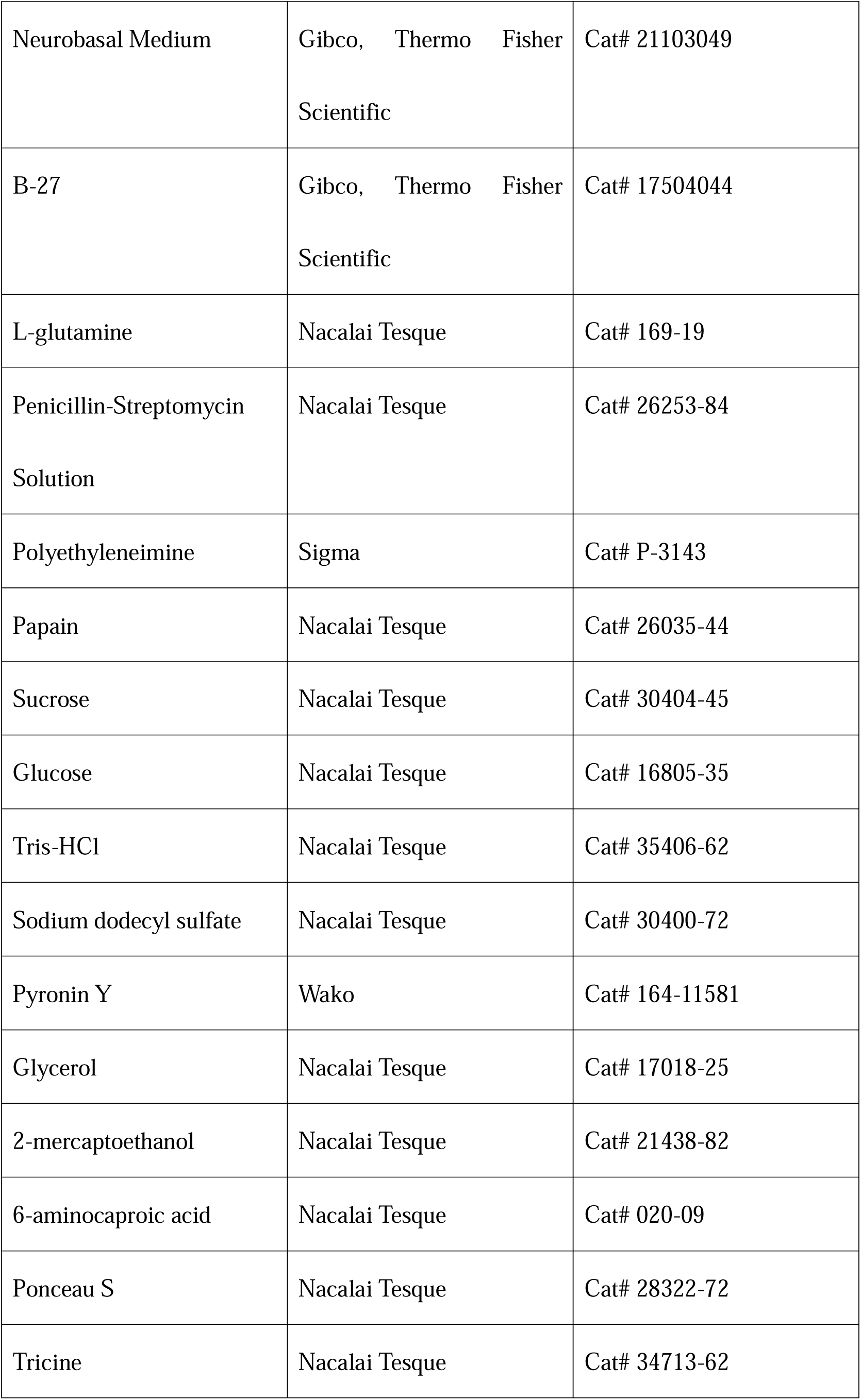
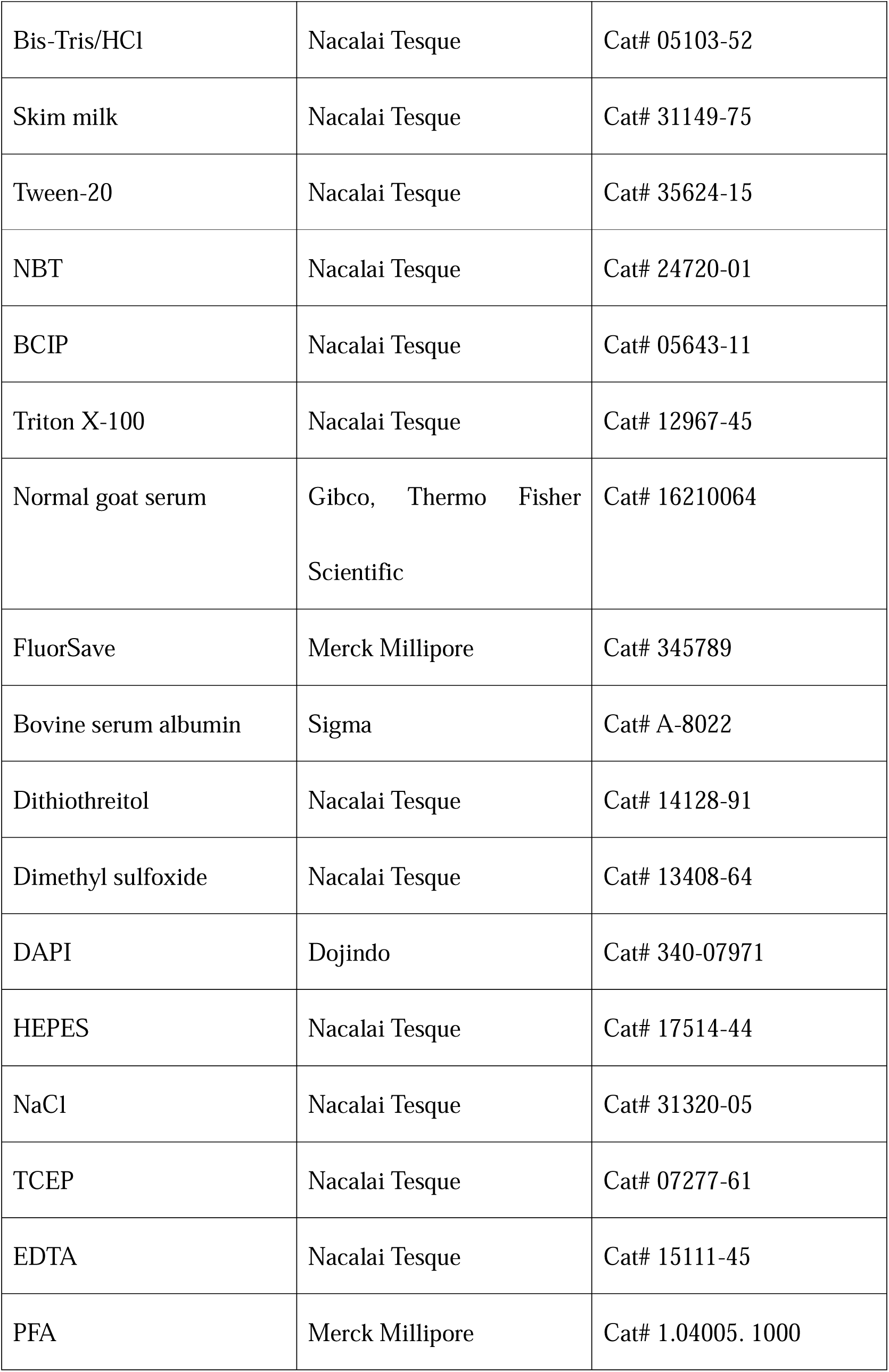
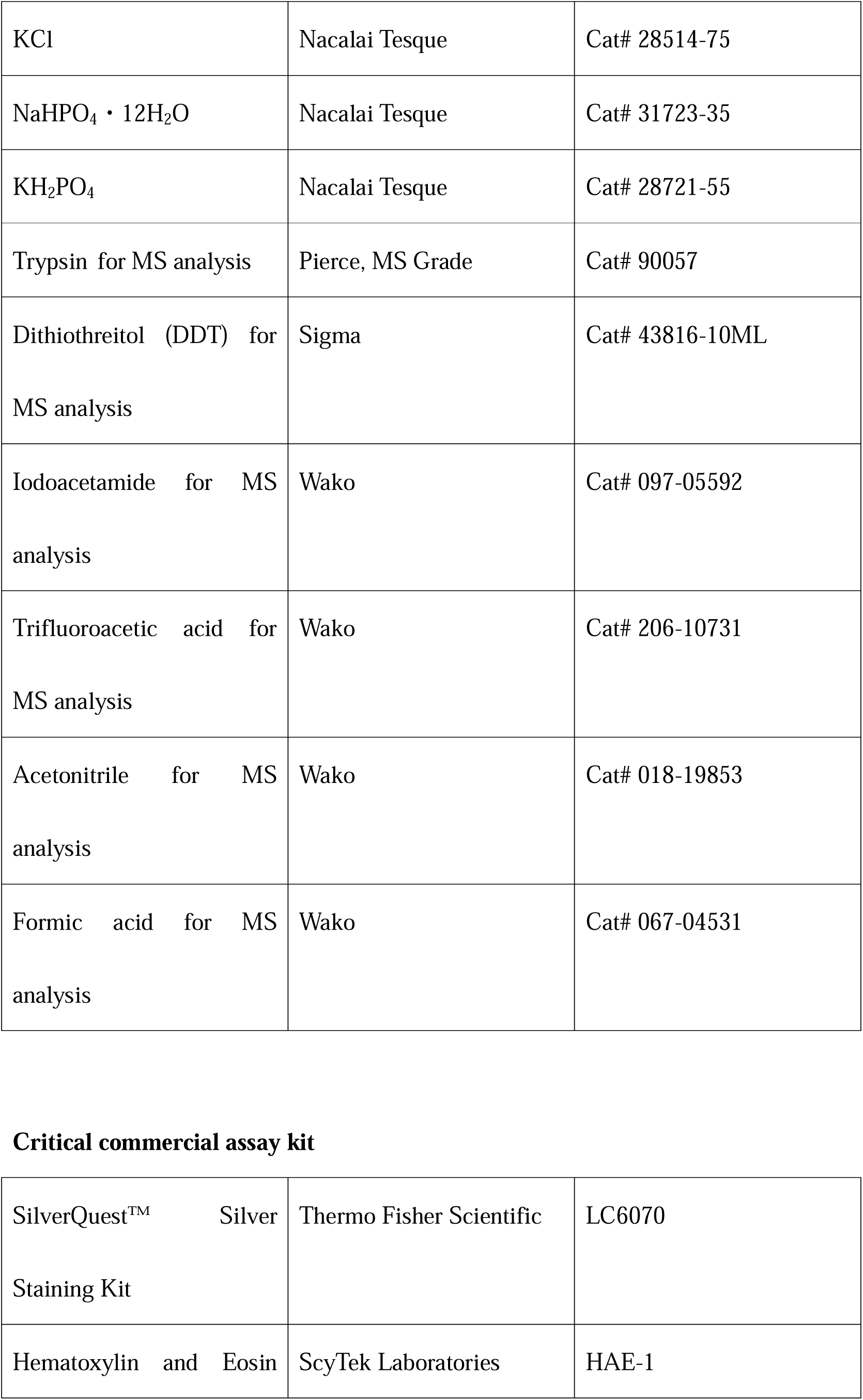
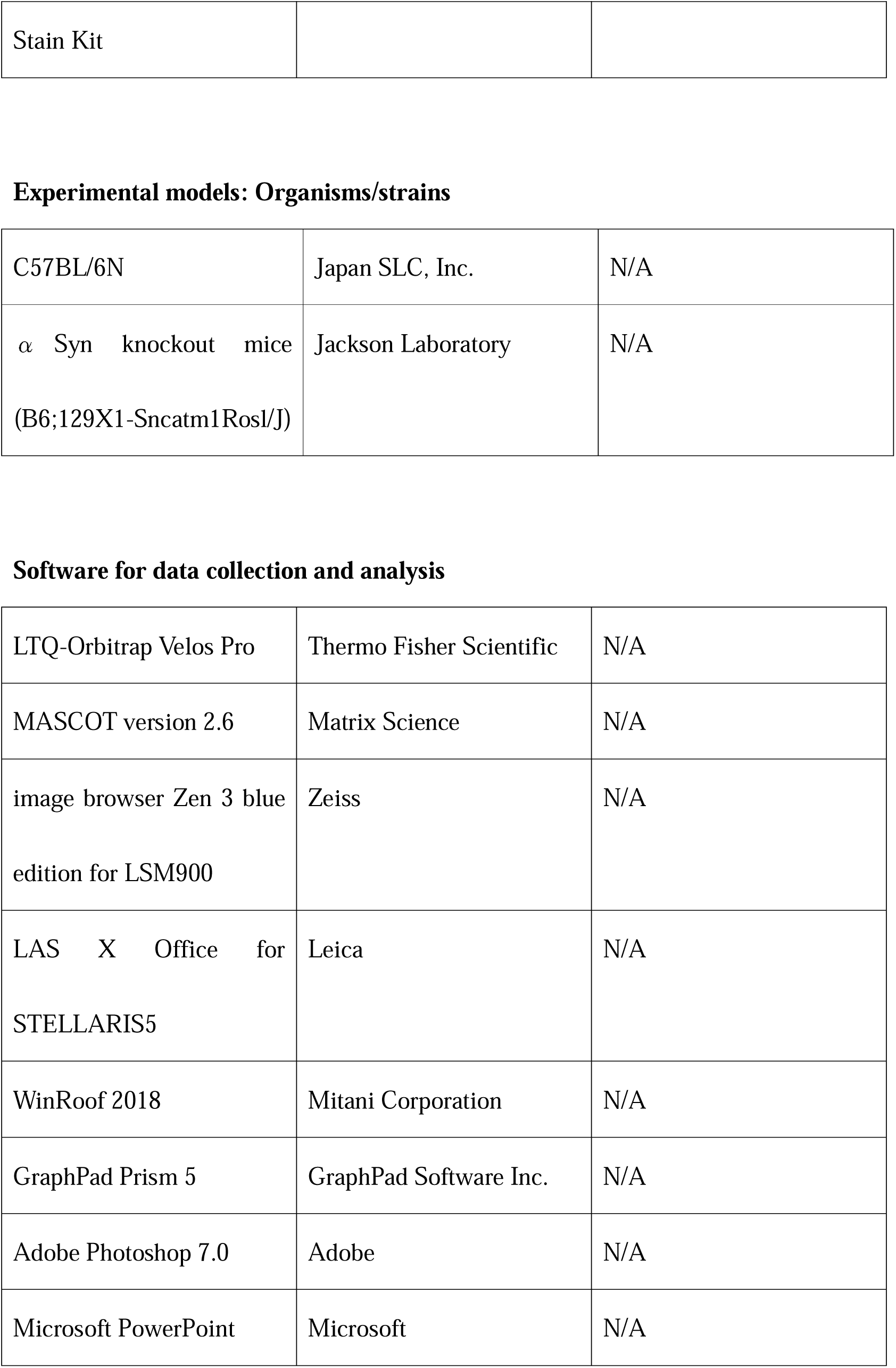

### Resource availability Lead contact

Further information and requests for resources and reagents as well as datasets and protocols used in this study should be directed to and will be fulfilled by the lead contact, KT and MT, as described below.

**Katsutoshi Taguchi, Ph.D.**

Department of Anatomy and Neurobiology, Graduate School of Medical Science, Kyoto Prefectural University of Medicine, Kawaramachi-Hirokoji, Kamikyo-ku, Kyoto 602-8566, Japan. Tel: +81-75-251-5301; Fax: +81-75-251-5306; AND

**Masaki Tanaka, M.D., Ph.D.**

Department of Anatomy and Neurobiology, Graduate School of Medical Science, Kyoto Prefectural University of Medicine, Kawaramachi-Hirokoji, Kamikyo-ku, Kyoto 602-8566, Japan. Tel: +81-75-251-5300; Fax: +81-75-251-5306;

## Materials availability

The experimental materials and data resources used in this study are available from the corresponding authors, MT and KT, upon reasonable request.

## Data and code availability

### Mass spectrometric and proteomic analyses

The raw files of the mass spectrometric analysis were used to conduct searches against the mouse and human αSyn amino acid sequences that were obtained from the Mus musculus dataset (Uniprot Proteome UP000000589 2021_03_07 downloaded), Homo sapiens dataset (Uniprot Proteome UP000005640 2022_05_25 downloaded), and the common Repository of Adventitious Proteins (cRAP; ftp://ftp.thegpm.org/fasta/cRAP). The search was conducted by using MASCOT version 2.6 (Matrix Science) via Proteome Discoverer 2.4 (Thermo Fisher Scientific).

Based on the results of mass spectrometric analysis, the candidate enzymes which process the N-terminal side of lysine residues located at the N-terminal repeat regions of αSyn were screened using the Peptidase database, MEROPS (https://www.ebi.ac.uk/merops/).

All the remaining research data are available in this article and its Supplementary Information. Furthermore, the source data have been provided with this paper.

## Experimental model and study participant details Animals

αSyn homozygous knockout (KO) mice (B6;129X1-*Snca^tm1Rosl^*/J^34^) were purchased from the Jackson Laboratory and maintained on a C57BL/6N background. Adult male C57BL/6N wildtype (WT) and αSyn-KO (12- to 18-week-old) mice were used in the histological experiments. KO mice were genotyped according to the manufacturer’s protocol (Jackson). The mice were housed under a 12-h light/dark cycle in a temperature-controlled environment with ad libitum food and water in our animal facility. All animal experimental procedures conformed to the Guidelines for Animal Experiments, Kyoto Prefectural University of Medicine (approval numbers: M2024-148, M2024-154, and M2024-509) and national regulations, and the number of animals used and their suffering were minimized.

## Ethics approval and consent to participate

The use of postmortem brain tissue in this study was approved by the Institutional Review Boards of Kyoto Prefectural University of Medicine (approval no. ERB-C-1846-2). Written informed consent for cadaver surgical training and the use of tissue samples for medical research was obtained, both from patients themselves and their next of kin.

## Consent for publication

Not applicable

## Method details

### Primary neuronal culture

Using a previously reported method^35^, with slight modifications, primary cultured neurons were obtained. Briefly, dissociated cells obtained from the prefrontal cortices and hippocampi of WT mouse embryos (E15–18) were disseminated onto polyethyleneimine-coated coverslips or into the chambers of microfluidic devices, and cultured in Neurobasal Medium (Gibco, Thermo Fisher Scientific) supplemented with B-27 (Gibco, Thermo Fisher Scientific), L-glutamine (Nacalai Tesque), and penicillin/streptomycin (Nacalai Tesque). Half of the volume of the culture medium was changed every 3–4 days until use. To induce the formation of intracellular phosphorylated αSyn aggregates, neurons were cultured in PFF-supplemented complete Neurobasal Medium (final 20 μg/mL) for 4–5 days, unless specified otherwise.

### Preparation of recombinant **α**Syn and PFFs

For the preparation of recombinant human αSyn from bacterial culture, plasmid vectors were constructed as described previously^36^. Briefly, PCR fragments of human αSyn were inserted into pTrc-His-TOPO vector (Thermo Fisher Scientific). His-tagged αSyn was purified by cOmplete™ His-Tag Purification Column (Roche). To confirm the purity of the protein by SDS-PAGE, the purified fraction was electrophoresed, and the protein band was stained by Coomassie Brilliant Blue. The final protein yield was determined with Lowry’s method.

PFFs were prepared according to a previously reported method, albeit with some slight modifications^11^. Briefly, PFFs were generated by incubating purified αSyn in sterile PBS (final concentration: 2 mg/ml) at 37°C with constant agitation for 5–6 days, then ultracentrifuging (at 200,000*g*) the PFF-containing solution for 2 h, and recovering the PFF precipitate at the bottom of the centrifuge tube. The resulting pellet was dissolved in sterile PBS and stored at −30°C until use.

To prepare recombinant αSyn harbouring mutations at the N-terminal repeat region (Figure 2D), a DNA fragment of mouse full-length αSyn harbouring those mutations was artificially synthesized and inserted into the pEX-A2J2 vector by Eurofins Genomics Japan. A DNA fragment of the mutated αSyn was further cloned into pTrc-His2B vector (Thermo Fisher Scientific). The presence of each mutation was confirmed by DNA sequencing using the ABI PRISM3130 Genetic analyzer. The His-tagged mutated αSyn-recombinant protein was purified using the abovementioned method described for WT αSyn.

### Microfluidic devices

Transmission of pathological αSyn was reproduced *in vitro* using microfluidic devices^37^. To isolate cell-to-cell transmissible pathological seeds, two types of microfluidic devices were used. For small-scale preparation of the seeds, three-compartment devices (TCND500) were purchased from Xona and cell culture was performed according to the manufacturer’s protocol. Briefly, pathological neurons harbouring intracellular phosphorylated-αSyn aggregates were cultured in the middle chamber, C2 (Figure 1A). Seeds released from those pathological neurons were recovered in the culture medium in the side chambers, C1 and C3, which are connected to C2 through microgrooves. During cell culture, to prevent the backflow of the PFF-containing medium into the C1 and C3 chambers, the medium in the C2 chamber was constantly maintained at a lower height than that of the C1 and C3 chambers. For large-scale preparation of seeds to perform mass spectrometric analysis, a large polydimethylsiloxane-based microfluidic device was developed (Figure S2A–C). Before the dissemination of dissociated cells prepared from the embryonic brain, the large microfluidic device was settled on a polyethyleneimine-coated glass dish that comprises two areas – a central square area and a hollow square (outer) area – which are connected by microgrooves through which the axons extend (Figure S2D, E). Pathological neurons were cultured in the central area of this device, and the seeds released from these neuronal cells were recovered from the medium in the outer area. During cell culture, the height of the medium in the central area was constantly maintained lower than that in the outer area to prevent the backflow of the PFF-containing medium.

### Preparation of the seed fraction

Unless specified otherwise, the seeds were prepared using the microfluidic devices. Partly seeds fraction was prepared using the batch method (Figure S1A–F). After the PFF treatment on primary neurons cultured in the middle or central square chamber, half of the volume of the seed-containing medium in the side (C1 and C3) or outer chambers was recovered every 3–4 days for 14 days. By ultrafiltration through a 3-kDa molecular-weight cutoff membrane (Merck Millipore), this seed-containing medium was concentrated up to 10 folds. Next, a discontinuous sucrose density gradient was prepared in a centrifuge tube purchased from Beckman Coulter (Figure 1C and Figure S1A). Each layer comprised phosphate-buffered saline (PBS, pH 7.4) containing the indicated concentration of sucrose. The concentrated medium was overlaid on a 0.8 M sucrose-containing PBS layer, ultracentrifuged at 200,000g for 12 h at 4°C, and an equal volume (250-300 μL) was aliquoted and further subjected to biochemical analysis. Seeds were recovered on the interface between the concentrated medium and the 0.8 M sucrose-containing PBS layer (Fraction 4 in Figure 1C).

For seed preparation using the batch method, the conditioned medium of pathological neurons was recovered as the seed-containing medium. After 2-day culture under PFF treatment, the PFF-containing medium was discarded, the cells were washed 4 times with complete Neurobasal Medium and then cultured in a fresh Neurobasal Medium for 5–7 days. This conditioned medium was collected and concentrated using ultrafiltration as described above. The concentrated medium was fractionated using sucrose density-gradient ultracentrifugation as well as the microfluidic method. Seeds were recovered at the interface between the concentrated medium and 0.8 M sucrose-containing PBS and subjected to biochemical analysis or aggregate-formation assay (Figure S1).

### Immunoblotting assay

For SDS-PAGE, samples were denatured by heating with 1% 2-mercaptoethanol-containing SDS-sample buffer (62.5 mM Tris-HCl, pH 6.8, 2% sodium dodecyl sulfate, 0.003% Pyronin Y, and 10% glycerol) at 98°C for 3 min. Proteins were separated by 15% polyacrylamide gel (Wako) and transferred to a polyvinylidene difluoride (PVDF) membrane (Bio-Rad Laboratories). For native-PAGE, the samples were treated with sample buffer (15% glycerol, 0.5 M 6-aminocaproic acid, 0.001% Ponceau S, and 50 mM Bis-Tris/HCl, pH 7.0). Proteins were separated by NativePAGE™ Bis-Tris Gel purchased from Thermo Fisher Scientific and electrophoresed using anode (50 mM Bis-Tris/HCl, pH 7.0) and cathode (50 mM Tricine, 15 mM Bis-Tris/HCl, pH 7.0) buffers. After electrophoresis, polyacrylamide gel was immersed in a transfer buffer containing 1% SDS, and the separated protein bands were also denatured in this step. The denatured proteins were transferred to the PVDF membrane; the membrane was blocked with 5% skim milk in Tris-buffered saline (TBS, pH 7.4) containing 0.1% Tween-20 for 1 h at room temperature. After treatment with primary antibody, the membrane was incubated with an alkaline phosphatase-conjugated secondary antibody. Protein bands were detected using the NBT-BCIP system (Nacalai Tesque). For αSyn detection, mouse monoclonal antibody Syn-1 (1:1000; BD Biosciences) or rabbit polyclonal antibody C-20 (1:2000; Santa Cruz Biotechnology) was used.

### Immunocytochemistry

Cultured cells were fixed with 2% paraformaldehyde (PFA) in the culture medium for 10 min at room temperature, permeabilized with 0.1% Triton X-100 in PBS for 10 min, and blocked with 10% normal goat serum (NGS) in PBS for 30 min. Next, cells were incubated for 1–2 h with primary antibodies in the blocking solution, washed with PBS, and treated with secondary antibodies. The primary antibodies were detected with Alexa488- and Alexa594-conjugated secondary antibodies (Thermo Fisher Scientific). After labelling with the secondary antibodies, cells were washed with PBS and then with milliQ water (Merck Millipore), before being mounted with FluorSave (Merck Millipore).

For αSyn detection, Syn-1 or C-20 antibodies were used. Phosphorylated αSyn was detected by mouse monoclonal antibody pSyn#64 (1:1000; Wako) or rabbit polyclonal antibody phospho-S129 αSyn (1:1000; Abcam). SENP2 was detected by rabbit polyclonal antibody (1:250; GeneTex). Low-magnification images were acquired by using immunofluorescent signals captured through a UPlanFl 10×/0.30 Ph1 objective with an inverted fluorescent microscope IX71 (Olympus) equipped with a cooled charge-coupled device camera (Olympus). For confocal observation, images were acquired as Z stacks (5–15 z-sections, 1-μm apart, 1024 × 1024 pixels) using a Plan-Apochromat 63×/1.40 Oil DIC objective (Carl Zeiss) with an inverted laser-scanning confocal microscope, LSM900 (Carl Zeiss) or using an HC PL APO CS2 63×/1.40 OIL objective (Leica) with a laser-scanning confocal microscope, STELLARIS5 (Leica) for high-resolution imaging (Figure 4A).

### Atomic force microscopy

To acquire images of the seed structure, the seed fraction was dropped onto the surface of a freshly cleaved mica plate (NILACO) and incubated for 5 min. Images were acquired using atomic force microscopy, SPM-9700 (SHIMADZU), operating in Dynamic Mode. The scanning parameters were as follows: operating point, 0.146–0.166 V; drive (tapping) frequency, 323–325 kHz; and scan rate, 1.0 Hz.

### Size-exclusion chromatography

To determine the molecular weight of the seeds, size-exclusion chromatography (SEC)–high-performance liquid chromatography-based analysis was performed using the SMART system (Pharmacia). Briefly, 50 μL seed fraction was injected into a sample tube and separated using the Superose 6 column comprising filtered PBS (pH 7.4) as the mobile phase. Each sample was eluted at a 40 μL/min flow rate; each 80 μL was collected as a fraction. Protein elution was monitored by UV absorbance at 280 nm. To obtain a standard curve for the molecular-weight determination, gel-filtration standards (Gel Filtration Calibration Kit HMW) purchased from GE Healthcare were separated using the same SEC column and flow rate. To detect the presence of the seed structure recovered in these SEC fractions, each SEC fraction was further observed using immunoelectron microscopy.

### Immunoelectron microscopy

For immunolabelling of seeds or PFFs, the seed fraction or sonicated PFFs were dropped onto the surface of formvar-coated molybdenum grids, PVF-M15 STEM Mo150P (Okenshoji), and incubated for 10 min. After incubation, the excess solution on each grid was discarded; the grid surface was further blocked with 1% bovine serum albumin (BSA) in PBS for 15 min and incubated with primary antibodies for 1 h. After washing in PBS, primary antibodies were detected with 10-nm gold (BBI Solutions) and/or 6-nm gold (Abcam)-conjugated secondary antibodies in PBS containing 1% BSA. After labelling, the grids were washed with PBS, stained with uranyl acetate, and observed using a JEM-1220 electron microscope (JEOL) at 80 keV.

For αSyn detection, the C-20 antibody was used as a pan-αSyn antibody. For the detection of oligomeric species of αSyn, the O2 (1:500; BioLegend) and 5G4 (1:500; Merck Millipore) antibodies, which recognize soluble oligomers and fibril forms of αSyn, including LBs and LNs^28,29^, were used.

### Mass spectrometric analysis for screening of the **α**Syn-cleavage site

To detect the cleavage sites of the seed-constituting αSyn from the mass spectrometric analysis sample, the seeds were concentrated by immunoprecipitation. Briefly, the seed fraction was prepared using large microfluidic devices as the original fraction for immunoprecipitation. The seeds were concentrated by immunoprecipitation after labelling by a C-20 antibody and precipitation with protein A-conjugated magnet beads (Bio-Rad), and then eluted by the SDS sample buffer at 56°C for 15 min. The eluted proteins were further denatured by heating at 98°C with 1% 2-mercaptoethanol for 3 min, and separated by SDS-PAGE. Protein bands were detected by silver staining using the SilverQuest™ Silver Staining Kit (Thermo Fisher Scientific). The PFF-constituting αSyn was prepared as the control for mass spectrometric analysis, because the results of the artificially cleaved sites detected in the PFF-constituting αSyn were subtracted from those of the cleaved sites detected in the seed-constituting αSyn. These protein bands (Nos. 1–5 and control bands, Figure 2B) were subjected to mass spectrometric analysis.

For mass spectrometric analysis using Nano-liquid chromatography–tandem mass spectrometry (LC–MS/MS), the proteins present in each gel slice underwent a series of stepwise treatment as follows: reduction with 10 mM dithiothreitol (DDT) at 56°C for 1 h, alkylation with 55 mM iodoacetamide at room temperature for 45 min in the dark, and digestion with 10 μg/mL modified trypsin (Pierce, MS Grade) at 37°C for 16 h. The resulting peptides were extracted using 1% trifluoroacetic acid and 50% acetonitrile, vacuum-dried, and dissolved in a solution of 2% acetonitrile and 0.1% formic acid.

Mass spectra were acquired using an LTQ-Orbitrap Velos Pro (Thermo Fisher Scientific) coupled with a nanoflow UHPLC system (ADVANCE UHPLC; AMR Inc.) featuring Advanced Captive Spray SOURCE (AMR Inc.). The peptide mixtures were loaded onto a C18 ID 0.1 mm × 20 mm, 5-μm (particle size) trap column (Acclaim PepMap 100 C18, Thermo Fisher Scientific) and fractionated through a C18, ID 0.075 × 500 mm, particle size 3-μm column (CERI). The peptides were eluted at a flow rate of 300 nL/min with a linear gradient of 5–35% solvent B over 60 min. Buffers A and B comprised 0.1% formic acid and 100% acetonitrile, respectively. The mass spectrometer was programmed for 11 successive scans, including a full-scan MS over 350–2,000 m/z by orbitrap at a resolution of 60,000, and automatic data-dependent MS/MS scans of the 10 most intense ion signals in the first precursor scan by ion-trap detectors for the second and eleventh scans. Using a 2 m/z isolation width and a 90-s exclusion time for molecules in the same m/z range, MS/MS spectra were obtained with a normalized collision energy of 35% CID.

The raw files were searched against the mouse and human αSyn amino acid sequences, including the N-terminal His-tag sequence, with Mus musculus dataset (Uniprot Proteome UP000000589 2021_03_07 downloaded), Homo sapiens dataset (Uniprot Proteome UP000005640 2022_05_25 downloaded), and the common Repository of Adventitious Proteins (cRAP; ftp://ftp.thegpm.org/fasta/cRAP). The search was conducted through MASCOT version 2.6 (Matrix Science) via Proteome Discoverer 2.4 (Thermo Fisher Scientific). The following settings were used as the search parameters: Enzyme, Semi-trypsin; false discovery rate (FDR), 0.01; precursor mass tolerance, 10 ppm; product tolerance, 0.8 Da; fixed modification, carbamidomethylation of cysteine; variable modification, oxidation of methionine, and acetylation of protein N-termini; and maximum missed cleavages, 2.

### Screening of the **α**Syn-processing enzyme and inhibitor assay

Based on the results of mass spectrometric analysis, the candidate enzymes which cleave the N-terminal side of lysine residues located at the N-terminal repeat regions of αSyn (conserved cleavage sites in Figure 2C) were screened using the Peptidase database, MEROPS (https://www.ebi.ac.uk/merops/); next, inhibitors for each enzyme were selected (Table 1), dissolved with dimethyl sulfoxide (DMSO), and stored at −30°C until use. To ascertain the inhibitory effect of the inhibitors on aggregate formation, primary neurons were cultured and treated with PFFs in the presence of each inhibitor. The following inhibitors were purchased from the manufacturers and used at the indicated final concentration: Mergepta (46 μM; MuseChem), carboxypeptidase inhibitor derived from potato tuber (CPI) (1 μM; Sigma), and NSC 632839 (9 μM; Selleckchem). As a control, primary neurons were cultured with diluted DMSO as well as each inhibitor.

### Image analysis and quantification of the inhibitory effect of NSC632839 on aggregate formation in primary cultured neurons

The phosphorylated αSyn-positive and DAPI-positive areas were measured using WinRoof 2018 software (Mitani Corporation). As shown in Figure S3, the phosphorylated-αSyn immunoreactive area, excluding the DAPI-positive area, was quantitatively analysed because the monoclonal antibody against phosphorylated αSyn #64 tends to react with nuclei. In either the presence or absence of NSC632839, wherein the threshold intensity was kept constant and set to include measurement profiles by visual inspections. Data represent the mean ± SEM. Statistical analysis was performed using GraphPad Prism 5 with a two-tailed, unpaired *t*-test and assessed using a significance level of *p*[<[0.05 (n=3 fluorescent images acquired from 2 coverslips were analysed for each treatment; two independent cultures were performed, and reproducibility was confirmed).

### Protease assay

To examine whether αSyn is processed by the several candidate enzymes summarized in Table 1, a protease assay was performed using recombinant active enzymes: 7 μg soluble or PFF forms of recombinant αSyn (substrate) was mixed with 0.5–5.2 μg of each recombinant enzyme in the following buffer to a final volume of 100 μL: USP7 (50 mM HEPES (pH8.0), 0.1 M NaCl, and 1 mM TCEP), SENP2 (50 mM HEPES, pH 8.0; 0.15 M NaCl; and 1 mM DTT), USP2 and UCHL1 (50 mM HEPES, pH 8.0, 0.15 M NaCl, 0.1 mM EDTA, and 1 mM DTT). PFFs were sonicated using the Hielscher ultrasonic processor UP50H with sonotrode MS1 (10 s on–10 s off, repeated 10 times on ice, at 100% amplitude setting) prior to use. Each substrate was incubated with each enzyme at 37°C for the indicated duration (4–24 h). The reaction was stopped by the addition of SDS sample buffer, and the samples were immediately heated with SDS-sample buffer at 98°C for 3 min and subjected to immunoblotting assay. Recombinant enzymes were purchased from the following manufacturers: human USP2 catalytic domain, CF (R&D Systems), human His6-USP7 Protein, CF (R&D Systems), GST-tagged human SENP2 catalytic domain (Enzo), and mouse active UCHL1 (ProSpec).

### Human PD brain

The use of postmortem brain tissue in this study was approved by the Institutional Review Boards of Kyoto Prefectural University of Medicine (approval no. ERB-C-1846-2). Written informed consent for cadaver surgical training and the use of tissue samples for medical research was obtained, both from patients themselves and their next of kin. The postmortem brain specimen was obtained from a patient with PD (age: 85 years, female), fixed using Thiel’s method, with slight modifications. Briefly, a fixative containing 5% formalin was perfused from the superior sagittal sinus. The brain was removed from the skull, immersed in 3.7% formalin for 14 days, and dissected into tissue blocks, including the substantia nigra (SN). Tissue blocks were further fixed with 4% PFA in 0.1 M phosphate buffer (PB) for 2 days. After fixation, these tissue blocks were subjected to immunohistochemistry as described below.

### PFF injection and inhibitor infusion *in vivo*

To reproduce the propagation of LB-like pathology *in vivo*, WT mice were anesthetized using a midazolam/medetomidine/butorphanol cocktail (4, 0.3, and 5 mg/kg, respectively), and 2 μL PFFs (7 mg/mL) was stereotaxically injected into the striatum, as reported previously^8^. A needle inserted into the left forebrain (+0.98 mm relative to Bregma, +1.70 mm from midline) was used to direct the inoculum into the striatum (+3.5 mm beneath the dura) and was injected using a 10-µL Hamilton syringe at a flow rate of 0.2 µL/min (total 14 μg PFFs per site), with the needle left in place for >20 min.

The inhibitor (NSC 632839, final concentration 73.5 μM in sterile saline) was infused into the left lateral ventricle using an Alzet osmotic pump (Model 2006) connected to a brain infusion kit (Type 3) in accordance with the manufacturer’s instructions (Alzet) for controlled-release of the reagent (0.15 µL/h for 42 days). Subjects were monitored regularly following their postoperative recovery, and sacrificed at 6 weeks after the surgery. The efficiency of the aggregate formation and inhibitory effect of NSC632839 on aggregate formation was determined using immunohistochemistry. Data from NSC635839-infused mice (n=5) and DMSO-infused control mice (n=3) were included in the statistical analysis as described below.

To study the effect of endogenous αSyn expression on aggregate formation, we injected PFF into the striatum of WT and αSyn-KO mice (Figure S6), and the PFF injections were performed as described above.

### Immunohistochemistry and Hematoxylin and Eosin (HE) staining

Unless specifically mentioned, a standard immunohistological procedure was performed on free-floating sections prepared from mouse and human brains.

For mouse brain, immunostaining was performed as previously described^38^. Briefly, under deep anesthesia, adult male mice were intracardially perfused with PBS followed by 4% PFA in PBS. Brains were dissected into blocks and postfixed using the same fixative for 12 h at 4°C. Coronal sections (40 μm) were obtained using a vibratome (Dosaka-EM), permeabilized with 0.3% Triton X-100 in PBS (PBST) for 1 h, and blocked with 10% NGS in PBST for 20 h. These sections were then incubated with the primary antibody in the blocking solution for 20 h, washed with PBST, and treated with a secondary antibody. For double immunostaining, this staining procedure was repeated. For the detection of phosphorylated αSyn, two types of antibodies were used: monoclonal mouse IgG antibody (#64) and polyclonal rabbit IgG antibody (pS129 αSyn).

For the immunofluorescent method, the primary antibodies were detected with Alexa488-and Alexa594-conjugated secondary antibodies (Thermo Fischer Scientific). Sections were washed with PBST and then with 20 mM Tris-HCl buffer, and mounted with FluorSave (Merck Millipore). For confocal observation, images were acquired as Z stacks (10–20 z-sections, 1-μm apart, 1024 × 1024 pixels) using a Plan-Apochromat 63x/1.40 Oil DIC objective (Carl Zeiss) with an inverted laser-scanning confocal microscope, LSM900 (Carl Zeiss).

For the immunoenzyme method, the primary antibody was detected by biotinylated anti-rabbit IgG antibody (1:500; Vector Laboratory) for 4 h, followed by horseradish peroxidase (HRP)-conjugated streptavidin (1:2; Nichirei Biosciences Inc.) detection for 1 h. HRP was visualized with 3,3’-diaminobezidine tetrahydrochloride and H_2_O_2_ in 50 mM Tris-buffered saline (pH 7.4). Immunosignals were acquired through the UPlanFl 60× objective with an inverted microscope (IX71), equipped with a cooled charge-coupled device camera (Olympus).

For the human PD brain, the dissected tissue block including the SN was cryoprotected using 20% sucrose in 0.1 M PB (pH 7.4) and rapidly frozen in Tissue-Tek OCT compound (Sakura Finetek Japan). Sections were obtained using a cryostat (Leica) at 30-μm thickness. For HE staining, sections were stained on a slide glass using a Hematoxylin and Eosin Stain Kit according to the manufacturer’s standard protocol (ScyTek Laboratories). For immunostaining with antigen retrieval, sections were immersed in formic acid for 10 min, washed 4 times with PBS, and subjected to immunohistochemical analysis using anti-phosphorylated αSyn (#64) and anti-SENP2 antibodies with the immunostaining procedure described for mouse brain sections.

### Quantification of the number of cell bodies harbouring pathological aggregates of phosphorylated **α**Syn

The number of cell bodies harbouring aggregates of phosphorylated αSyn in the amygdala and piriform cortex of NSC632839-treated and DMSO-treated control mouse brains was counted. For cell-counting in each region, 3–6 sections per animal for each treatment were subjected to the cell-counting procedure. The number of animals used in each experimental group is indicated in the figure legends (Figure 5F, G). After the cell-counting procedure, the mean cell numbers per animal were individually calculated in each brain region; the mean values per experimental group were then used in the statistical analysis using GraphPad Prism 5 by the two-tailed, unpaired *t*-test, with or without Welch’s correction. Data are expressed as the mean ± SEM. The significance levels were set at \**p* < 0.05.

### Image processing

The digital images obtained on the LSM900 or IX71 were processed using Adobe Photoshop 7.0 (Adobe). Each figure set was prepared in Microsoft PowerPoint (Microsoft).

### Statistical analysis and reproducibility

Statistical analyses between the two experimental groups were performed using the two-tailed, unpaired *t*-test, with or without Welch’s correction (GraphPad Prism 5, GraphPad Software Inc.). The number of animals used in each experiment and the number of images that were analysed are specified in the Methods and/or indicated in each figure legend. Data are expressed as the mean ± standard error of the mean. The significance levels were set at *p* < 0.05 (*) and *p* < 0.01 (**), as indicated. Reproducibility was confirmed by independent experiments that were repeated at least two to three times, with similar results.

## Acknowledgments

We thank Professor Masaya Ikegawa (Genomics, Proteomics and Biomedical Functions, Department of Life and Medical Systems, Faculty of Life and Medical Sciences, Doshisha University, Kyoto, Japan) for facilitating the pilot experiments in mass spectrometric analysis and thank Dr. Reiko Nakagawa (Laboratory for Cell-Free Protein Synthesis, RIKEN Center for Biosystems Dynamics Research, Kobe, Japan) for the mass spectrometric analysis and the preparation of the datasheets. This work was supported by Grants-in-Aid for Scientific Research from the Japan Society for the Promotion of Science (KT: 24K10514, 21K07299, and 18K07371; YW: 24K10494; MT: 21K06412) and The Shimizu Foundation for Immunology and Neuroscience Grant 2021 (to KT).

## Author contributions

KT and MT conceived and designed the experiments. KT performed the experiments. YW contributed to the analysis and discussion. KT and MT wrote the manuscript.

## Declaration of interests

The authors have no potential or actual conflicts of interest.

## Supplemental information

### Supplemental data figures and legends

**Figure S1.**
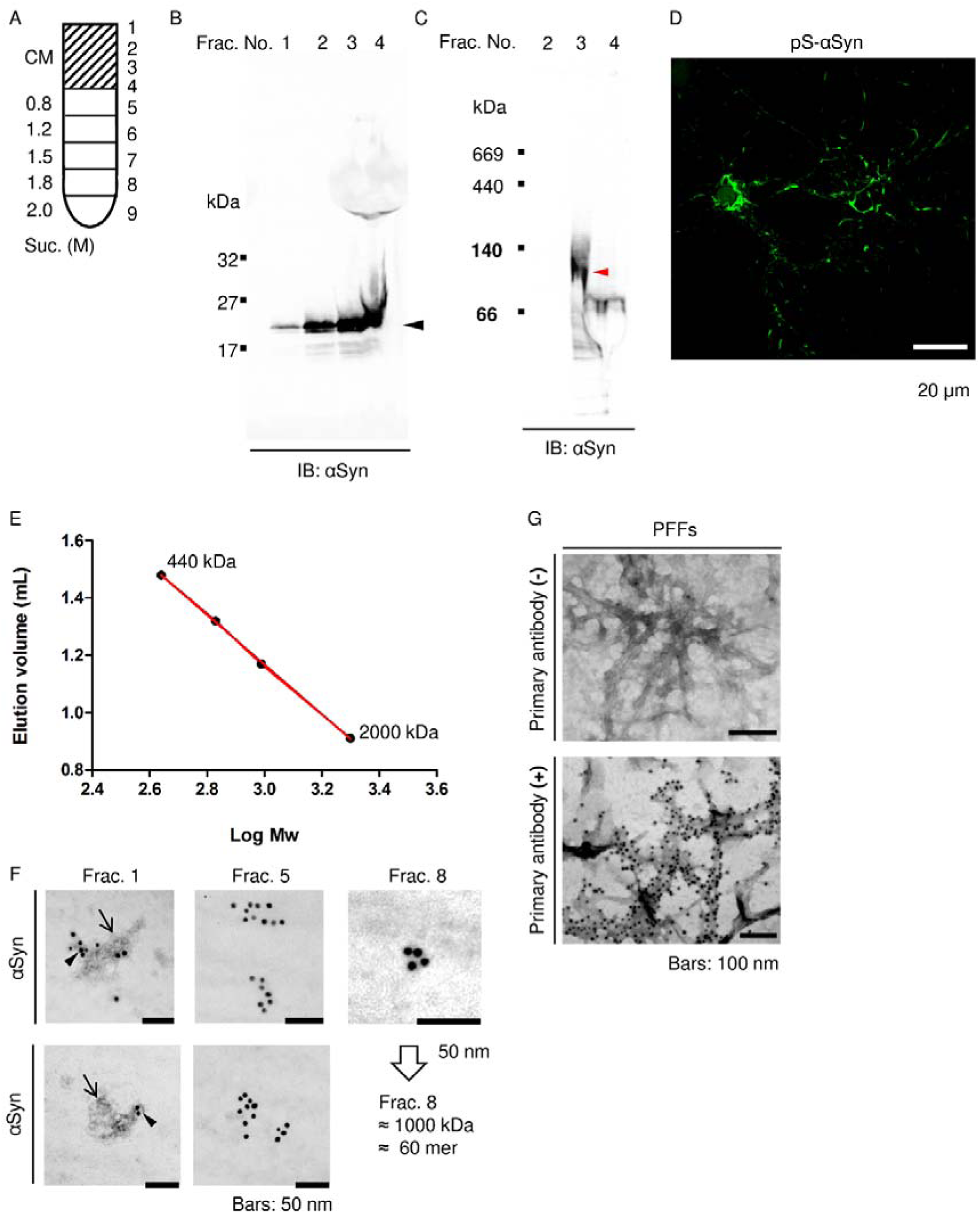
Isolation and characterization of pathological seeds using the batch method. **(A)** Schematic diagram of fractionation by sucrose-density gradient ultracentrifugation of the conditioned culture medium derived from pathological primary neurons harbouring Lewy body- and Lewy neurite-like aggregates. CM, culture medium. **(B)** SDS-PAGE and an immunoblotting assay of the fractions using anti-αSyn antibody. The numbers indicate the recovered fractions shown in **(A)**. **(C)** Native-PAGE and immunoblotting assay of the fractions. **(D)** The seeding ability of the pathological seeds recovered in Fraction 3. pS-αSyn, phosphorylated αSyn. **(E)** Standard curve for HPLC (Log. molecular weight vs. elution volume). **(F)** Immunoelectron microscopic observation of the αSyn-oligomer species recovered in each HPLC-prepared fraction. **(G)** Immunoelectron microscopic observation of PFFs as a control for αSyn immunolabeling. PFFs, preformed fibrils. Scale bar: 20 μm in **(D)**, 50 nm in **(F)**, and 100 nm in **(G)**.

**Figure S2.**
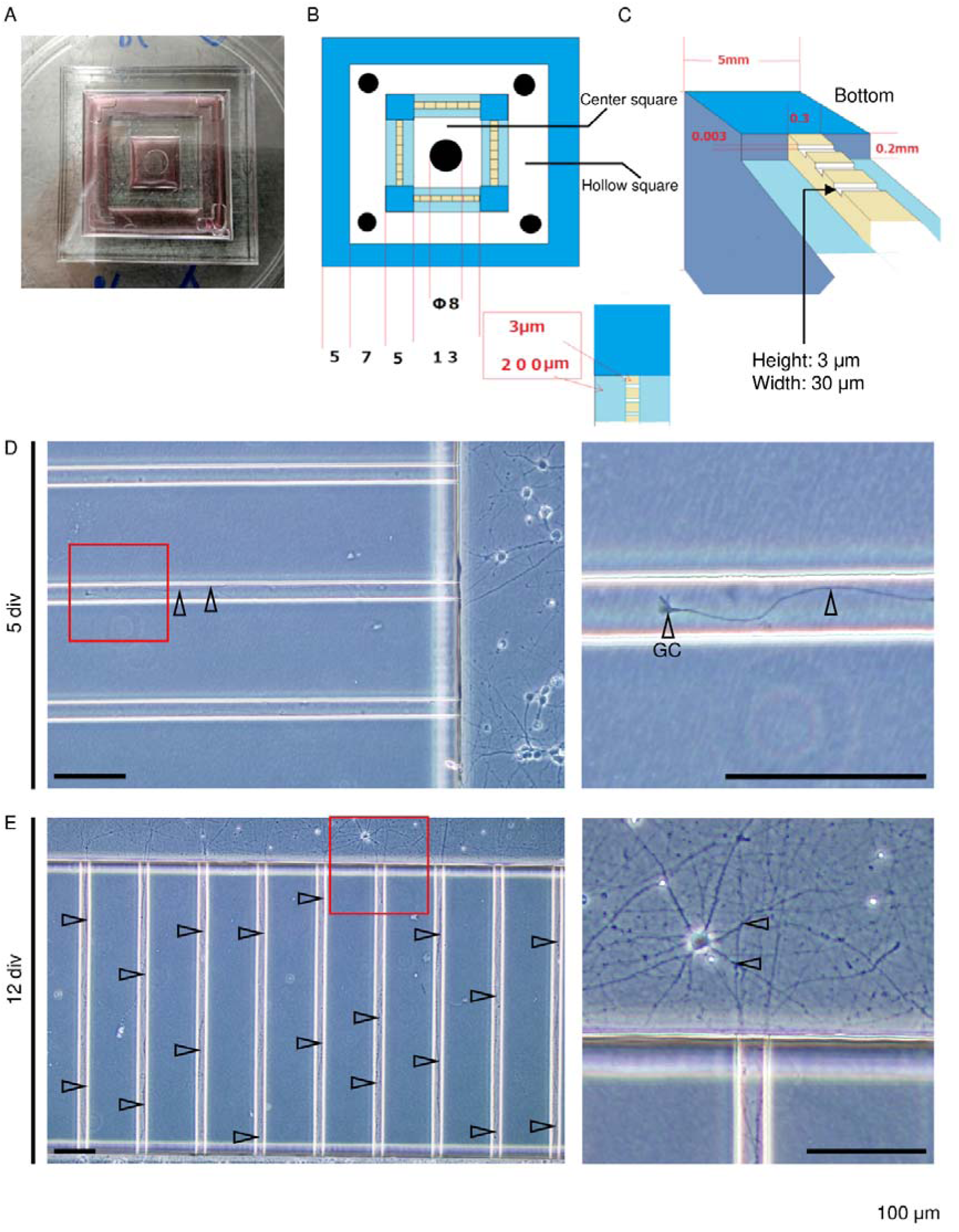
Development of the microfluidic device for large-scale preparation of pathological seeds. **(A–C)** Overview of microfluidic device for large-scale preparation of seeds. **(D, E)** Primary neuronal culture using the large-scale microfluidic device for 5 days **(D)** and 12 days **(E)** *in vitro*. Arrowheads show the extending axons. The region marked by a red square in **(D)** and **(E)** is magnified in each right panel, respectively. GC, growth cone. Scale bar: 100 μm.

**Figure S3.**
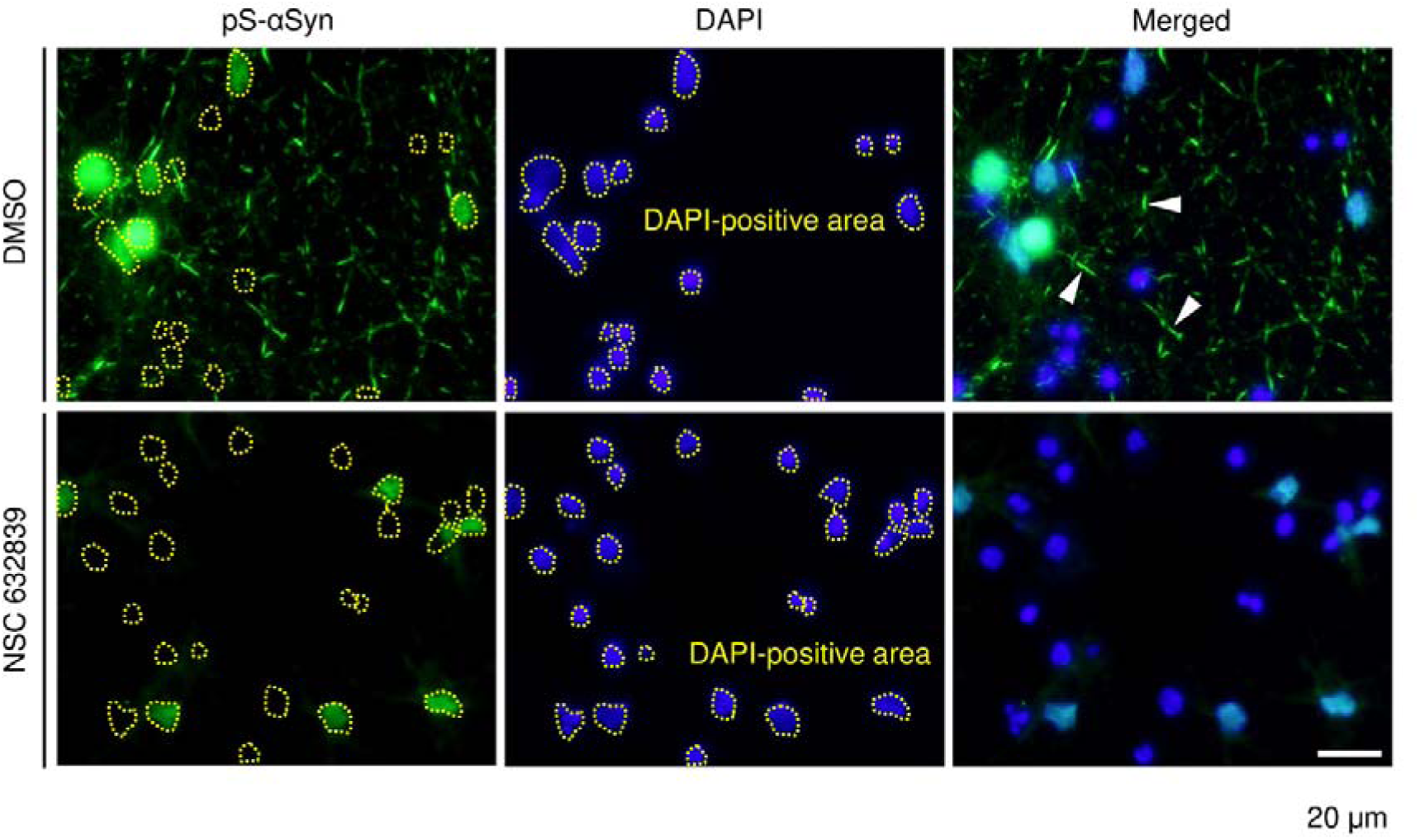
Image analysis of the inhibitory effect of NSC632839 on aggregate formation. Phosphorylated-αSyn immunoreactive area, excluding the DAPI-positive area surrounded by dotted lines, was quantitatively analysed because the monoclonal antibody against phosphorylated αSyn #64 tends to react with the nuclei. Arrowheads indicate examples of phosphorylated αSyn-positive aggregates. Scale bar: 20 μm.

**Figure S4.**
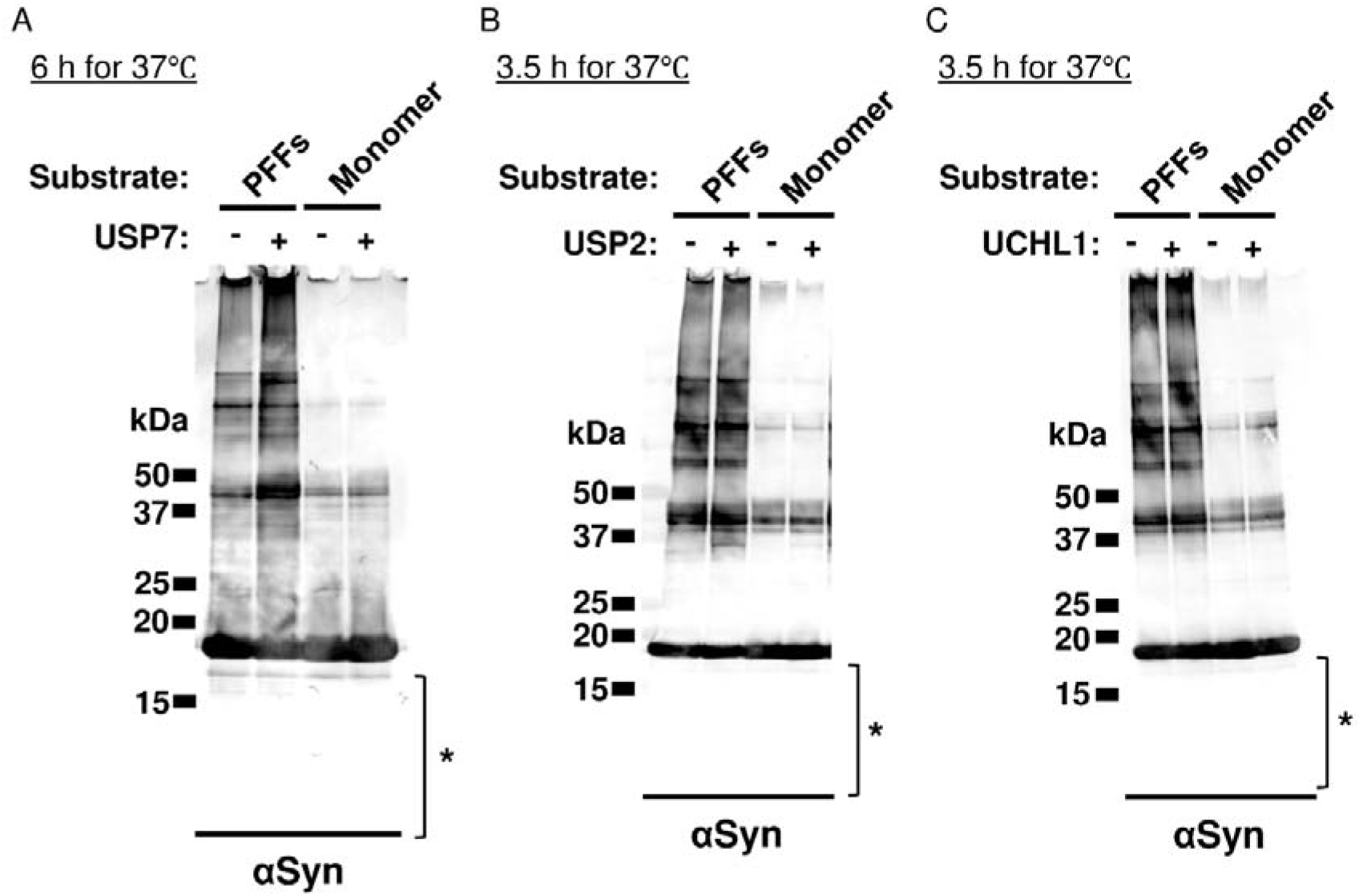
Screening assay for possible αSyn-processing enzymes. **(A–C)** αSyn-processing ability of possible NSC632839-inhibited enzymes was examined by a protease assay using recombinant proteins. As indicated by asterisks, cleaved fragments were not detected in the presence of USP-7 **(A)**, USP-2 **(B)**, or UCHL1 **(C)**. Both PFFs and a monomeric form of αSyn were not processed as appropriate substrates by these enzymes. The reaction time and temperature are indicated in each figure panel.

**Figure S5.**
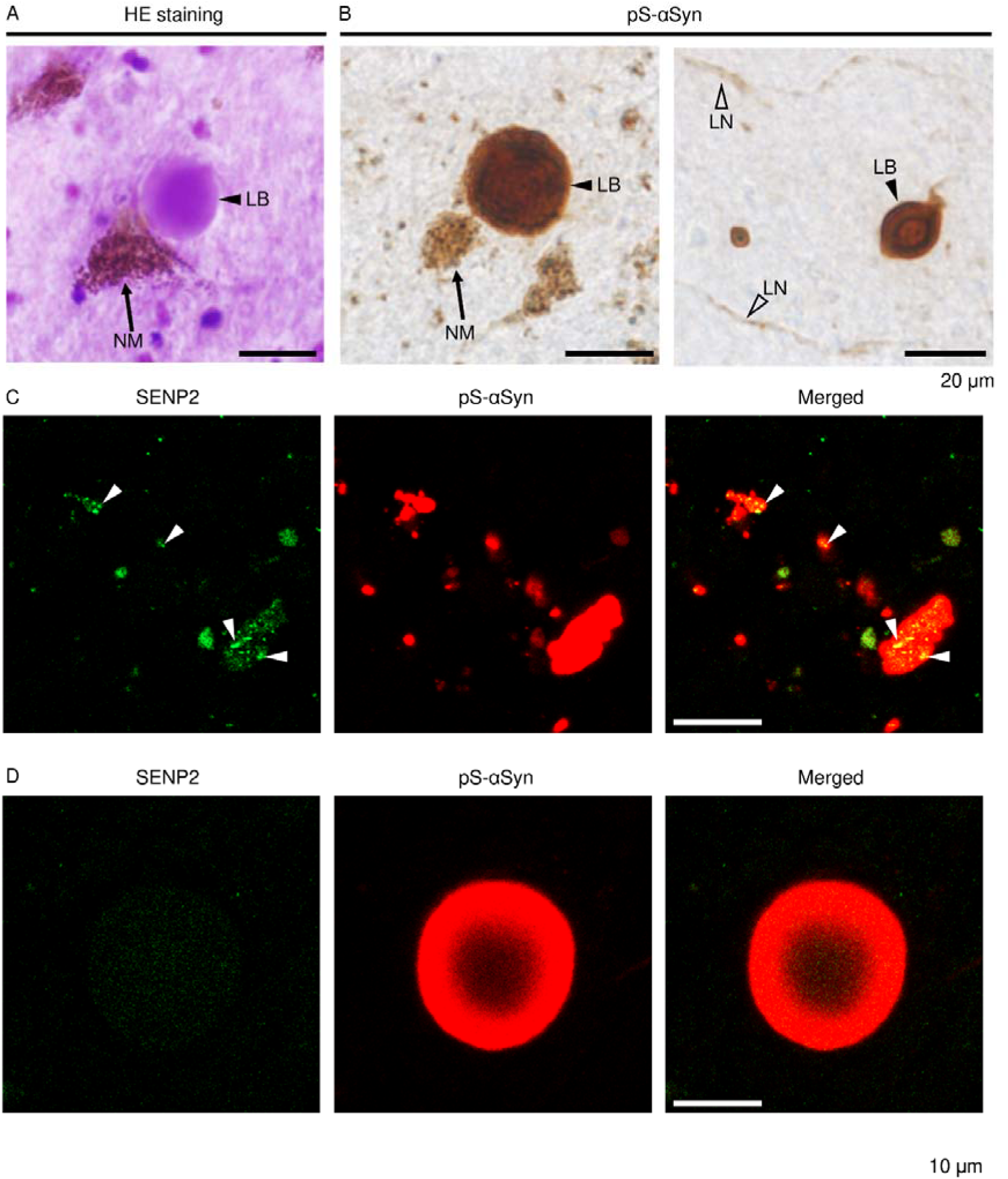
Presence of SENP2 in Lewy pathology. **(A**, **B)** Lewy body, Lewy neurites, and neuromelanin observed in the substantia nigra pars compacta of the PD-brain. **(C, D)** Presence of endogenous SENP2 in amorphous aggregates **(C)**, but not in a typical Lewy body **(D)**. Arrowheads indicate the presence of both SENP2 and phosphorylated αSyn. LB, Lewy body; LN, Lewy neurites; NM, neuromelanin. Scale bar: 20 μm in **(A, B)**, and 10 μm in **(C, D)**.

**Figure S6.**
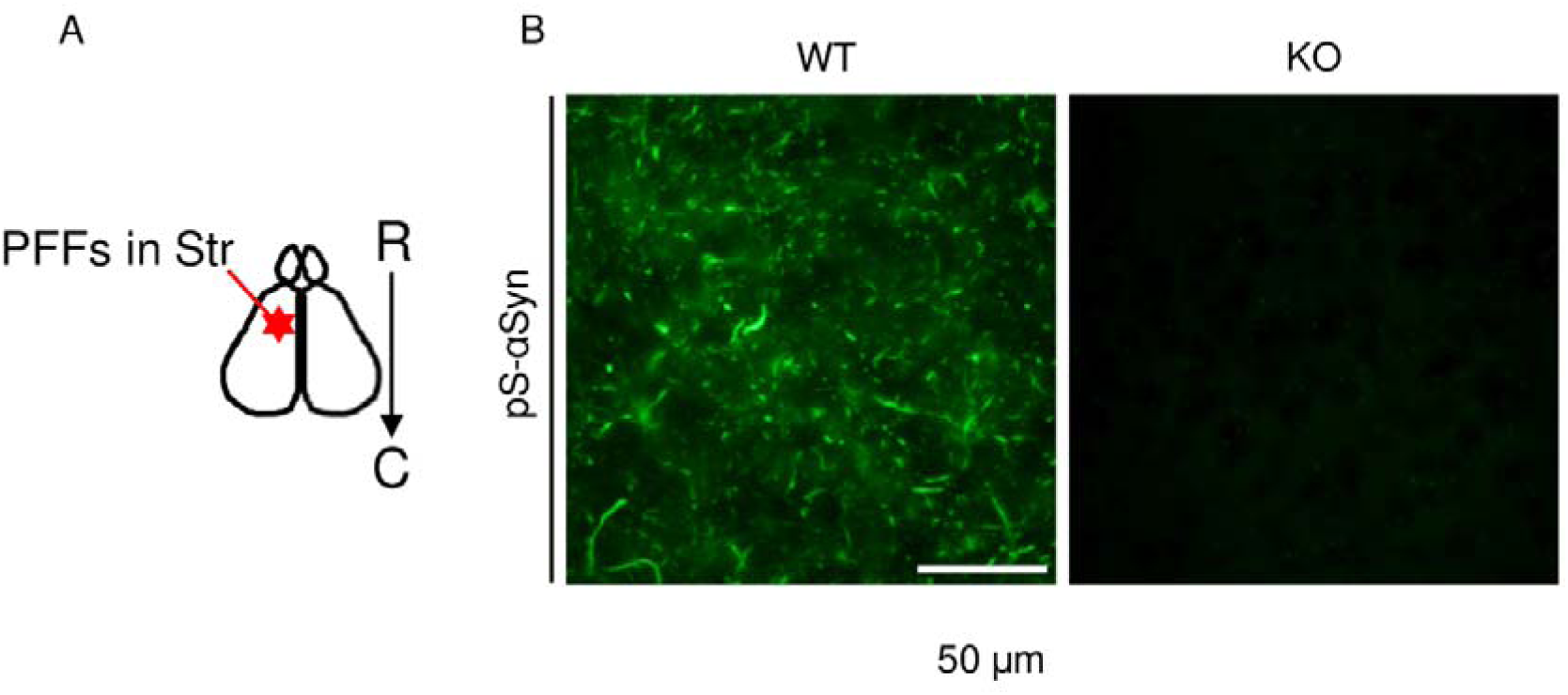
Endogenous αSyn expression is indispensable for aggregate formation. **(A)** Schematic diagram of PFF injection into the striatum of the mouse brain. C, caudal; R, rostral; Str, striatum. **(B)** Intense formation of phosphorylated-αSyn aggregates was observed in the WT, but not αSyn-KO, mouse brain. WT, wildtype; KO, knockout. Scale bar: 50 μm.

## Notes

### Competing Interest Statement

The authors have declared no competing interest.

### Summary of Updates

The format of main text was slightly modified.

